# An *SMN2* splicing-correcting antisense oligonucleotide promotes gene looping and transcription activation

**DOI:** 10.1101/2025.10.10.681673

**Authors:** Jose N. Stigliano, Emilia Haberfeld, Luciano E. Marasco, Ana Fiszbein, Nick J. Proudfoot, Alberto R. Kornblihtt

**Affiliations:** Universidad de Buenos Aires (UBA), Facultad de Ciencias Exactas y Naturales, Departamento de Fisiología, Biología Molecular y Celular and CONICET-UBA, Instituto de Fisiología, Biología Molecular y Neurociencias (IFIBYNE), 1428 Buenos Aires, Argentina; Sir William Dunn School of Pathology, University of Oxford, Oxford OX13RE, United Kingdom; Biology Department, Boston University, Boston, 02215, USA

## Abstract

We have previously proposed a combined therapy for spinal muscular atrophy based on the upregulation of exon 7 (E7) inclusion into *SMN2* mRNA by the antisense oligonucleotide nusinersen (ASO1) together with histone deacetylase inhibitors (HDACi). Using a CRISPR/deadCas-based strategy, we show here that targeting histone acetylation at the *SMN2* E7 region duplicates the HDACi effect, which overcomes pleiotropic effects. Most noticeably, targeted acetylation at the E7 region causes acetylation and activation of the 30-kbp-distant promoter. Since E7 and the nusinersen target site map near the *SMN2* gene 3’-end, this cross-talk between the two ends of the gene prompted us to use chromosome conformation analysis (3C) to find that ASO1 promotes gene looping. This unpredicted property of ASOs in modulating chromatin 3D configuration depends on cohesin, CTCF, cleavage/polyadenylation factors and a canonical poly(A) signal, and causes promoter activation at a long distance, which expands the therapeutic potential of splicing-correcting ASOs.

## INTRODUCTION

Loss-of-function mutations of the human *SMN1* gene are the cause of spinal muscular atrophy (SMA). Humans have a paralog, *SMN2*, whose exon 7 (E7), located towards the 3’ end of the gene, is mostly skipped. Consequently, it cannot fully compensate for the lack of *SMN1* expression. Both *SMN2* and *SMN1* differ in only 11 bp of their approximately 30-kbp lengths, which includes a C>T base change 6 bp inside exon 7 that causes E7 skipping^1^, giving rise to a truncated and unstable version of the SMN protein. One of the strategies that has been developed to treat SMA involves the splicing-correcting antisense oligonucleotide (ASO) nusinersen (a.k.a. Spinraza), that targets a splicing silencer located in *SMN2* intron 7 pre-mRNA and, by blocking the binding of the negative splicing factors hnRNPA1 and A2, it promotes higher E7 inclusion, increasing full length SMN protein levels^2–4^.

According to the kinetic coupling model, slow transcriptional elongation can promote either exon inclusion or skipping, depending on the type of exon^5–7^. In class I exons, slow elongation promotes inclusion by improving the recruitment of constitutive or positive splicing factors to pre-mRNA, whereas in class II exons slow elongation enhances the binding of inclusion-inhibitory factors to their target sites, promoting exon skipping. Moreover, intragenic deployment of specific histone marks can inhibit or promote RNAPII elongation by affecting chromatin compaction, which consequently affects alternative splicing. For example, dimethylation of histone H3 lysine 9 (H3K9me2) promotes class I exon inclusion^8,9^, whereas histone acetylation along gene bodies promotes skipping of the same class of exons^10^. Conversely, intragenic histone acetylation upregulates class II exon inclusion, which is consistent with the fact that inhibition of elongation promotes class II exon skipping^6^.

We have recently demonstrated that *SMN2* E7 is a class II alternative exon, which allowed us to use kinetic coupling tools to promote its inclusion in combination with nusinersen^11^. We found that by promoting transcriptional elongation, histone deacetylase (HDAC) inhibitors such as valproic acid (VPA) cooperate with a nusinersen ASO (hereafter ASO1) to upregulate *SMN2* E7 inclusion. Surprisingly, ASO1 also elicits the deployment of the silencing histone mark H3K9me2 on the *SMN2* gene, creating a roadblock to RNAPII elongation that acts negatively on E7 inclusion. By removing the roadblock, HDAC inhibition counteracts the undesired chromatin effects of the ASO, resulting in higher E7 inclusion. These results not only allowed us to propose a combined therapy for SMA but raised novel mechanistic questions such as to what extent ASO-elicited histone methylation, and its balance with histone acetylation, is critical for the splicing-correcting activity of the nusinersen ASO. We therefore tested whether replacement of the genome-wide, potentially pleiotropic, effect of VPA by targeted histone acetylation at the *SMN2* gene was an effective strategy.

We show here that the ASO1-elicited deployment of H3K9me2 not only occurs at the ASO1 target region but also at the 30-kbp-distant promoter. Furthermore, using a CRISPR/Cas-based strategy with a non-cleaving Cas9 fused to the transcriptional activator VP64 (dCas9-VP64), we show that targeting histone acetylation at the *SMN2* E7 gene region enhances the E7 inclusion levels caused by ASO1, and in addition, and most noticeably, the 30-kbp-distant promoter results activated. Reciprocally, targeting histone acetylation at the promoter enhances upregulation of E7 inclusion by ASO1. This crosstalk between the E7 alternative splicing event, mapping near the 3’ end of the gene, and the promoter region prompted us to use chromosome conformation capture (3C) analysis which revealed that ASO1 promotes looping between the 3’ and 5’ ends of the *SMN2* gene. This unforeseen feature of ASOs to influence chromatin 3D configuration is independent from the promotion of H3K9 dimethylation, occurs with different ASO chemistries and cell types and causes promoter activation. Most importantly, depletion and mutation analyses show a major role of the pre-mRNA cleavage and polyadenylation machinery on *SMN* gene looping.

## RESULTS

### Technical disclaimer

It should be noted that the *SMN1* and *SMN2* genes are present within a 500 kb duplicated region on chromosome 5. Due to the extremely high sequence identity between them, reads of deep sequence analyses and primers used to amplify segments in, for example, ChIP-qPCR protocols (see below) cannot be assigned unequivocally to either *SMN* gene, and so an assessment in parallel is imposed. Alternatively, the specific mRNAs splicing products of the human endogenous *SMN2* gene can be identified in the end-point RT-PCRs because *SMN2* exon 8 contains a *DdeI* restriction site that is absent in *SMN1*. Hence, RT-PCR followed by restriction digestion with *DdeI* distinguishes *SMN2* and *SMN1* transcripts^12^ . We will refer to merged *SMN1/2* genes when a parallel assessment is performed and as *SMN2* alone when a gene-specific method is applied. It is worth-mentioning that the nusinersen ASO indistinctively binds to its target sequence present in both *SMN1* and *SMN2* pre-mRNAs and therefore the therapeutic treatment is effective on, but not exclusive to, the *SMN2* gene.

### ASO1-elicited H3K9 dimethylation modulates its splicing-correcting activity

We have previously shown^11^ that nusinersen (ASO1) has two opposite effects on *SMN2* E7 splicing. On the one hand, by displacing the negative splicing factors hnRNPA1/A2 from their target sites at *SMN2* pre-mRNA, it promotes E7 inclusion. On the other hand, by eliciting H3K9 dimethylation, it has the opposite effect of enhancing E7 skipping (Figure 1A, left). Promotion of global histone acetylation via histone deacetylase inhibition with trichostatin A (TSA) or valproic acid (VPA) counteracts the negative chromatin effect of ASO1 on E7 splicing (Figure 1A, right). Consistent with this model, both RNAi depletion (Figure S1A and B) of the histone-dimethylating enzymes G9a and GLP in HEK293T cells (Figure 1B) and knockout^13^ of the G9a gene (Figure 1C) potentiate the E7 inclusion activity of ASO1. Additional evidence supporting a role for the H3K9me2 mark is provided by the fact that RNAi depletion of HP1β (Figure S1D), a reader of the H3K9me2 mark, enhances the E7 promoting activity of ASO1 (Figure 1D).

**Figure 1.**
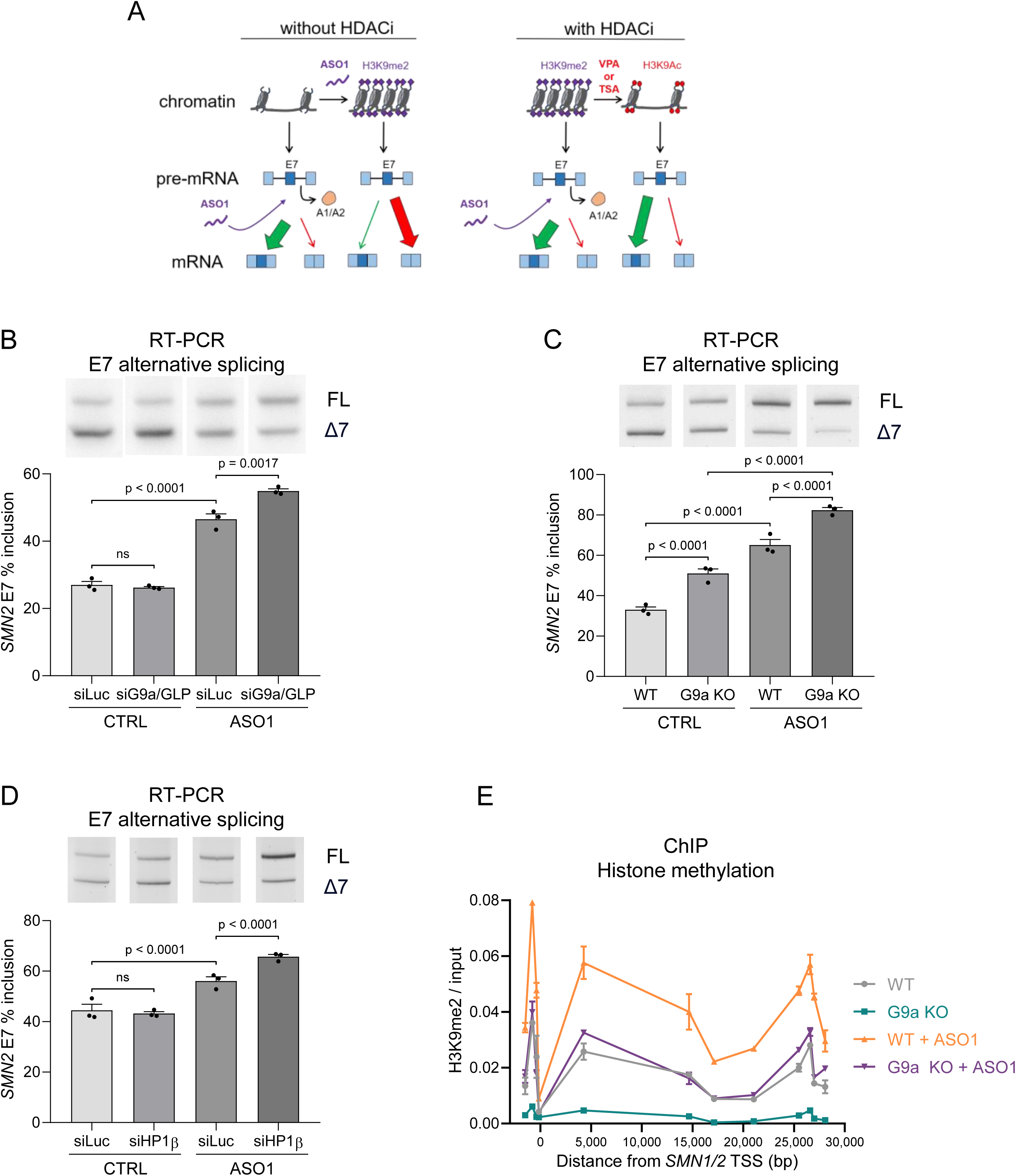
SMN2 exon 7 inclusion is regulated by H3K9 dimethylation. (A) Schematic model of ASO1 action on RNA and chromatin, adapted from Marasco et al. (2022). Without HDAC inhibition (left), ASO1 binding to *SMN2* pre-mRNA displaces hnRNP A1/A2 and promotes exon 7 (E7) inclusion, but concurrently induces H3K9me2 deposition, leading to chromatin compaction, reduced RNAPII elongation, and exon skipping. With HDAC inhibitors (right), histone acetylation counteracts chromatin compaction while ASO1 still enhances E7 inclusion, potentiating the splicing response. (B–D) RT-PCR analysis of *SMN2* E7 splicing in HEK293T cells. DdeI-digested PCR products resolved full-length (FL; upper band) and E7-skipping (Δ7; lower band) isoforms; representative gels (top) and E7 inclusion (%) from n = 3 biological replicates (bottom). Conditions: (B) siLuc or combined G9a/GLP knockdown (50 nM, 72 h), with or without ASO1 (1 nM, 72 h); (C) WT or G9a KO cells with control ASO or ASO1 (1 nM, 72 h); (D) siLuc or siHP1β (50 nM, 72 h), with or without ASO1 (1 nM, 72 h). (E) ChIP-qPCR analysis of H3K9me2 enrichment relative to input along the *SMN1/2* locus in WT and G9a KO HEK293T cells, with or without ASO1 (100 nM, 72 h). Interrogated regions are indicated as distance from the *SMN1/2* transcription start site (TSS). n = 2 biological replicates. Data are mean ± SEM. B–D, two-factor beta regression models followed by Tukey’s multiple-comparisons test. Significant p values are shown; ns, not significant; p < 0.05.

Chromatin immunoprecipitation (ChIP) analysis along the *SMN1/2* gene with an antibody to the H3K9me2 histone mark reveals that in G9a-KO cells, H3K9 dimethylation is practically absent along the entire gene compared with wild type cells (Figure 1E, blue and gray curves). ASO1 treatment increases H3K9 dimethylation both in wild type and G9-KO cells, but in the latter, ASO1 treatment is unable to reach the wild type levels (orange and purple curves). As far as the histone mark distribution elicited by ASO1 is concerned, both in WT and KO cells it peaks around the ASO1 target site, but also, and noticeably, at the promoter region, located approximately 30 kbp upstream.

Summarizing, the results in this section provide genetic evidence for the model of Figure 1A and reveal the specificity of H3K9me2, demonstrating the involvement of HP1β (Figure 1D). However, the fact that ASO1 also promotes histone methylation at the promoter (Figure 1E) suggests that the chromatin effects of ASO1 may have consequences other than those observed on splicing.

### Targeted histone acetylation counteracts the H3K9 methylation effect on splicing

Histone H3K9 methylation and several histone H3K acetylation marks a such as H3K9, K27, K18, K14, and K23 are usually considered to be antagonistic^14^. While H2K9me is a silencing mark, the mentioned H3K acetylation marks are usually permissive for transcription. Figure S1C confirms the antagonistic effects by showing that parallel to reduction in total H3K9 dimethylation there is a significant increase in total H3K9 acetylation in G9a-KO compared to WT cells. In our previous report^11^ we exploited the opposite effects of histone methylation and acetylation using histone deacetylase inhibitors (HDACi) in combination with ASO1, which supported the model in Figure 1A. However, while ASO1 acts specifically on the *SMN1/2* gene, histone deacetylase inhibitors act globally in the cell, which carries two challenges. On the therapeutic aspect, HDACi may have pleiotropic effects that could restrict their use in a combined therapy. At the mechanistic level, the effect of HDACi may be an indirect consequence of changes in expression of other genes or in acetylation of proteins other than histones. To overcome these obstacles, we set out to determine the effects of targeting histone acetylation specifically to the *SMN1/2* gene using a dCas9 (dead Cas9) strategy, in which a mutated, catalytically inactive, Cas9 enzyme is fused to the transcriptional activator VP64^15^. dCas9-VP64 acts by recruiting transcription machinery to target sites specified by CRISPR guide RNAs and is widely used to activate gene transcription upon promoter targeting in the so called CRISPR activation (CRISPRa) strategy. Since VP64 is known to recruit the histone acetyl transferase p300^16^, we used two sets of guide RNAs directed respectively to the E7 and the promoter regions of *SMN1/2* (Table S1, Figure 2A) to investigate their effects on ASO1-induced E7 inclusion. In a typical experiment, HEK293T cells were co-transfected with a plasmid vector expressing the dCas9-VP64 fusion protein and either a non-targeting guide RNA (NT) or guide RNAs targeting the relevant *SMN1/2* gene regions, with or without the ASO1 oligonucleotide. The NT guide RNA was designed to have a sequence with no complementarity in the human genome. Targeting dCas9-VP64 to the E7 region alone had no effect on basal E7 inclusion, but increased by approximately twofold the inclusion level elicited by ASO1 (Figure 2B). To test if its effect on ASO1-controlled splicing was in fact due to histone acetylation, we performed a similar experiment in the presence of the selective p300 inhibitor CBP30^17^, using the vehicle DMSO as control (Figures 2C and D). CBP30 abolishes the significant enhancement of E7 inclusion caused by ASO1. Consistent with the preference of p300 to acetylate histone H3K27^18^, western blot analysis indicates that the effect of CBP30 can be mostly attributed to inhibition of the H3K27ac mark (Figure S2).

**Figure 2.**
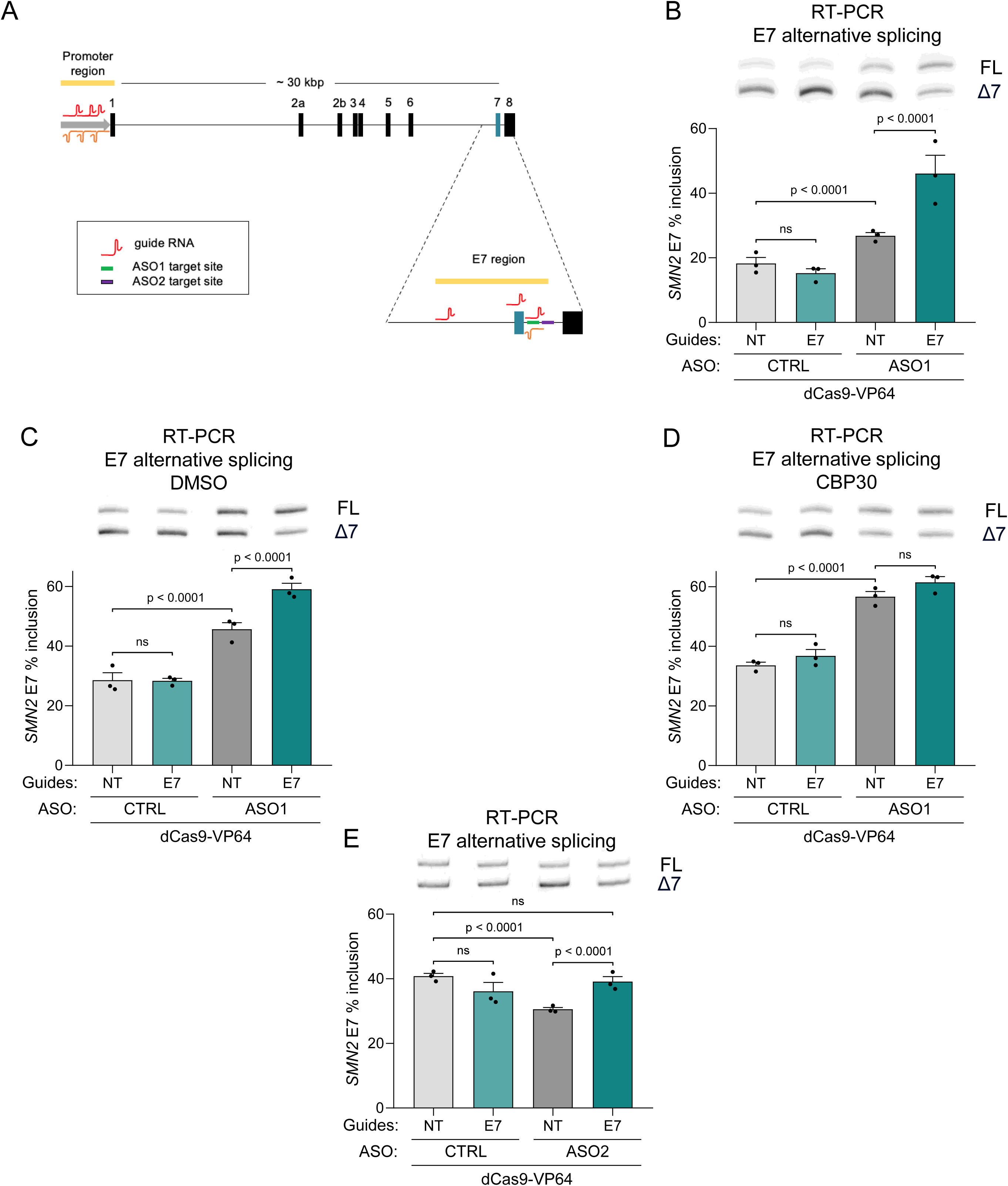
dCas9-VP64 targeting of the SMN2 E7 region synergizes with ASO1 to promote exon 7 inclusion. (A) Schematic of the *SMN2* locus and dCas9-VP64 targeting strategy. Black boxes, exons; yellow, promoter; green, ASO1 binding site; purple, ASO2 binding site; red (sense) and orange (antisense) wavy lines, designed sgRNA binding sites near promoter and E7 regions. (B–E) RT-PCR analysis of *SMN2* E7 splicing in HEK293T cells cotransfected with plasmids encoding dCas9-VP64 and non-targeting (NT) or E7-targeting sgRNAs (96 h). FL, full-length (upper band); Δ7, E7-skipping (lower band); representative gels (top) and E7 inclusion (%) from n = 3 biological replicates (bottom). Conditions: (B) WT cells with or without ASO1 (1 nM, 72 h); (C–D) with or without ASO1 (1 nM, 72 h) plus DMSO vehicle 0.2% (C) or selective p300/CBP inhibitor CBP30 (D; 40 µM, 48 h); (E) with or without ASO2 (50 nM, 72 h). Data are mean ± SEM. B–E, two-factor beta regression models followed by Tukey’s multiple-comparisons test. Significant p values are shown; ns, not significant; p < 0.05.

In our previous report we investigated the effects of a second ASO, ASO2, whose target site is also located in intron 7, but downstream of the ASO1 target site (Figure 2A). ASO2 bears no sequence identity with ASO1, and does not overlap hnRNPA1/A2 binding sites. Acting only at the chromatin but not at the pre-mRNA level, ASO2 causes H3K9 dimethylation both at the ASO target site and the gene promoter, resulting in a roadblock to RNAPII elongation at the E7 region and subsequent inhibition of *SMN2* E7 inclusion into mature mRNA^11^. We show now that targeting histone acetylation at the E7 region rescues the reduction E7 inclusion caused by ASO2 (Figure 2E).

In summary, specific acetylation at the E7 region acts similarly as global acetylation by HDAC inhibitors in counteracting the negative effects of both ASO1 and ASO2-mediated H3K9 dimethylation and therefore promotes E7 inclusion.

### Distant effects of targeted histone acetylation

As expected, ChIP analysis confirms that targeting dCas9-VP64 to the E7 region causes local histone acetylation both in the presence and in the absence of ASO1 (Figure 3A). Inspired by the long-distance effects of ASO1 in increasing histone methylation at the promoter (Figure 1E), we wanted to see if histone acetylation at the E7 region had a similar behavior. Figure 3B shows that, indeed, directing dCas9-VP64 to the E7 region also causes histone acetylation at the promoter region, even though this effect is only observed in the absence of ASO1. Consistently, this promoter histone acetylation is accompanied by transcription activation (Figure 3C) that also occurs when dCas9-VP64 is targeted to the promoter itself (Figure 3D), as expected for a classical CRISPRa. In view of these distant effects and taking into account old results from our group demonstrating that promoter identity and occupation regulate alternative splicing^19–23^, we investigated the effects of promoter acetylation on E7 alternative splicing. Notably, directing dCas9-VP64 to the promoter not only causes higher E7 inclusion *per se* but also enhances E7 inclusion levels elicited by ASO1 (Figure 3E).

**Figure 3.**
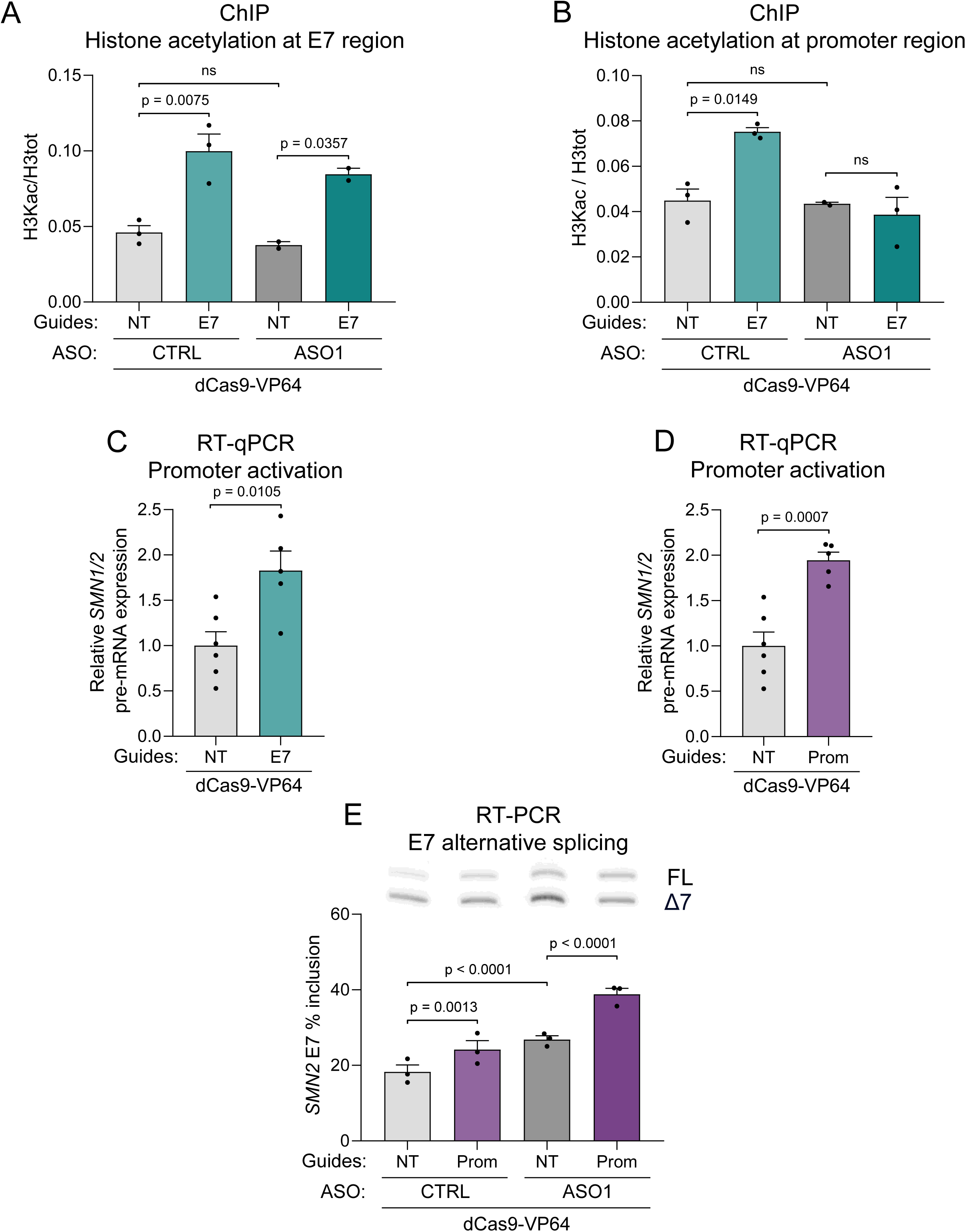
dCas9-VP64 induces histone acetylation and promoter activation at the SMN2 locus. HEK293T cells were cotransfected with plasmids encoding dCas9-VP64 and the indicated sgRNAs for 96 h. (A–B) ChIP-qPCR of pan-H3 acetylation (H3K9ac, H3K14ac, H3K18ac, H3K23ac, and H3K27ac), relative to total H3, at *SMN2* E7 (A) or promoter (B), using non-targeting (NT) or E7-targeting sgRNAs with control ASO or ASO1 (100 nM, 72 h). n = 2–3 biological replicates. (C–D) RT-qPCR of promoter activation using *SMN1/2* E1–I1 primers to detect pre-mRNA as a proxy for transcriptional output. (C) NT or E7-targeting sgRNAs; (D) promoter-targeting sgRNAs and the same NT control as in (C). n = 5–6 biological replicates. (E) RT-PCR of *SMN2* E7 splicing using NT or promoter-targeting sgRNAs, with or without ASO1 (1 nM, 72 h). FL, full-length (upper band); Δ7, E7-skipping (lower band); representative gel (top) and E7 inclusion (%) from n = 3 biological replicates (bottom). Data are mean ± SEM. Statistics: two-way ANOVA/Tukey (A–B), two-tailed Student’s *t* test (C–D), or two-factor beta regression/Tukey (E). Significant p values are shown; ns, not significant; p < 0.05.

These experiments allow us to conclude that localized histone acetylation, either at the promoter or at the E7 region, has conspicuous distant roles in the control of transcription and alternative splicing.

### ASOs promote gene looping

Results in Figures 1E, 3B, 3C and 3E demonstrate that there is a crosstalk between events occurring at the E7 region, located towards the 3’ end of the *SMN2* gene, and its promoter. To investigate if this was due to physical interactions between the 5’ and 3’ ends of the gene we performed chromosome conformation capture (3C)m analysis^24^. We designed sets of primers to assess the frequency of contacts between the promoter (anchor) and sequences at the 3’ end region or a transcriptional enhancer, reported approximately 1.8 kbp upstream of the +1 site of the *SMN1/2* gene merge^25^. Contacts between the anchor and intron 1 were used as a control. When cells were treated with ASO1, contacts between the promoter and I1 or between the promoter and the enhancer were not affected. However, importantly, there was a significant increase in the contact frequency between the promoter and the 3’ end region (Figure 4A).

**Figure 4.**
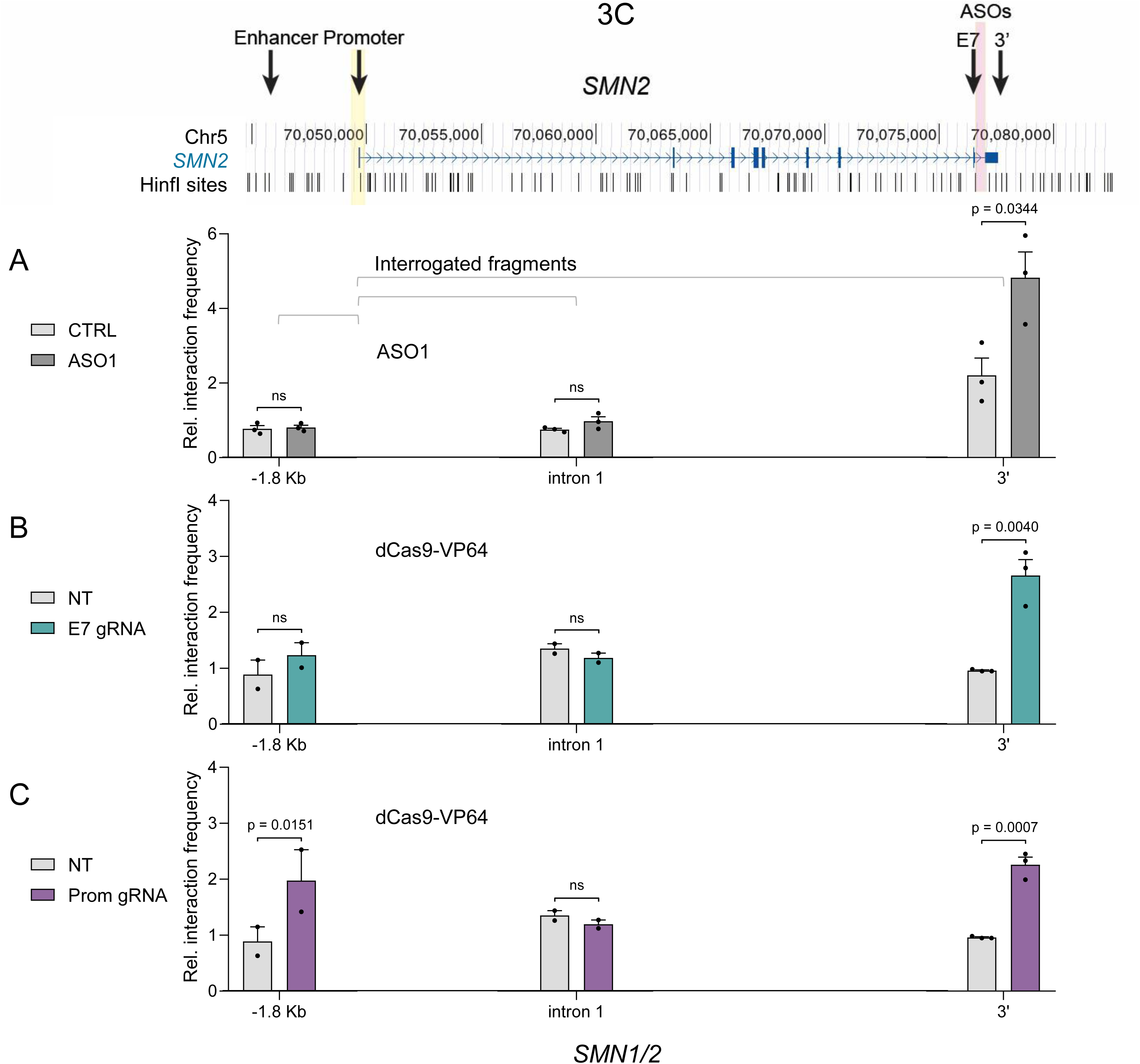
ASO1 and dCas9-VP64 promote *SMN2* gene looping. Chromatin conformation capture (3C)-qPCR analysis of *SMN1/2* chromatin interactions in HEK293T cells. Top, UCSC Genome Browser view (GRCh38/hg38) of *SMN2*, showing exons (blue boxes), introns (arrows), and HinfI sites. The anchor primer was immediately upstream of the TSS (yellow line); interactions were interrogated 1.8 kb upstream of the promoter (enhancer), within intron 1, and downstream of the 3′ end, and normalized to a U6 gene amplicon. The ASO1 target site is indicated by a pink line. (A) Cells with or without ASO1 (100 nM, 72 h; dark and light gray, respectively). (B– C) Cells cotransfected for 96 h with dCas9-VP64 and NT sgRNA (gray) or E7-targeting (B; blue) or promoter-targeting (C; purple) sgRNAs; the NT control in (C) is the same as in (B). Data are mean ± SEM from n = 2–3 technical replicates. Experiments were independently repeated where possible with consistent outcomes. Two-tailed *t* tests compared conditions within each region. Significant p values are shown; ns, not significant; p < 0.05.

### Targeted histone acetylation also promotes gene looping

In view of results in Figures 3C and E showing that targeted histone acetylation at the E7 region activates the promoter and that targeted acetylation at the promoter affects E7 alternative splicing, we decided to investigate if these treatments also promoted looping. Indeed, we see that directing dCas9-VP64 to the E7 region causes an increase in the contacts of the promoter with the 3’ end region, but not with the I1 and TE regions (Figure 4B). Consistently, directing dCas9-VP64 to the promoter increases its contacts with the 3’ end but not with the I1 region. However, unlike previous treatments in this figure, targeted histone acetylation at the promoter also increases its contacts with the transcriptional enhancer (Figure 4C), a phenomenon already observed for the effects of CRISPR-based transcriptional activation on enhancer-promoter interactions^26^.

### Both ASO1 and ASO2 activate the *SMN* promoter

We wanted to find out whether the looping effect was due to the pre-mRNA mechanism of ASO1 (the control of splicing by displacement of hnRNPA1/A2), or to its chromatin effects. To this end, we treated cells with ASO2, whose target site maps near that of ASO1 but that does not interfere with hnRNAPA1/A2 binding, and revealed that, as well as ASO1 (Figure 4A), ASO2 also promoted contacts of the promoter with the 3’ end region (Figure 5A).

**Figure 5.**
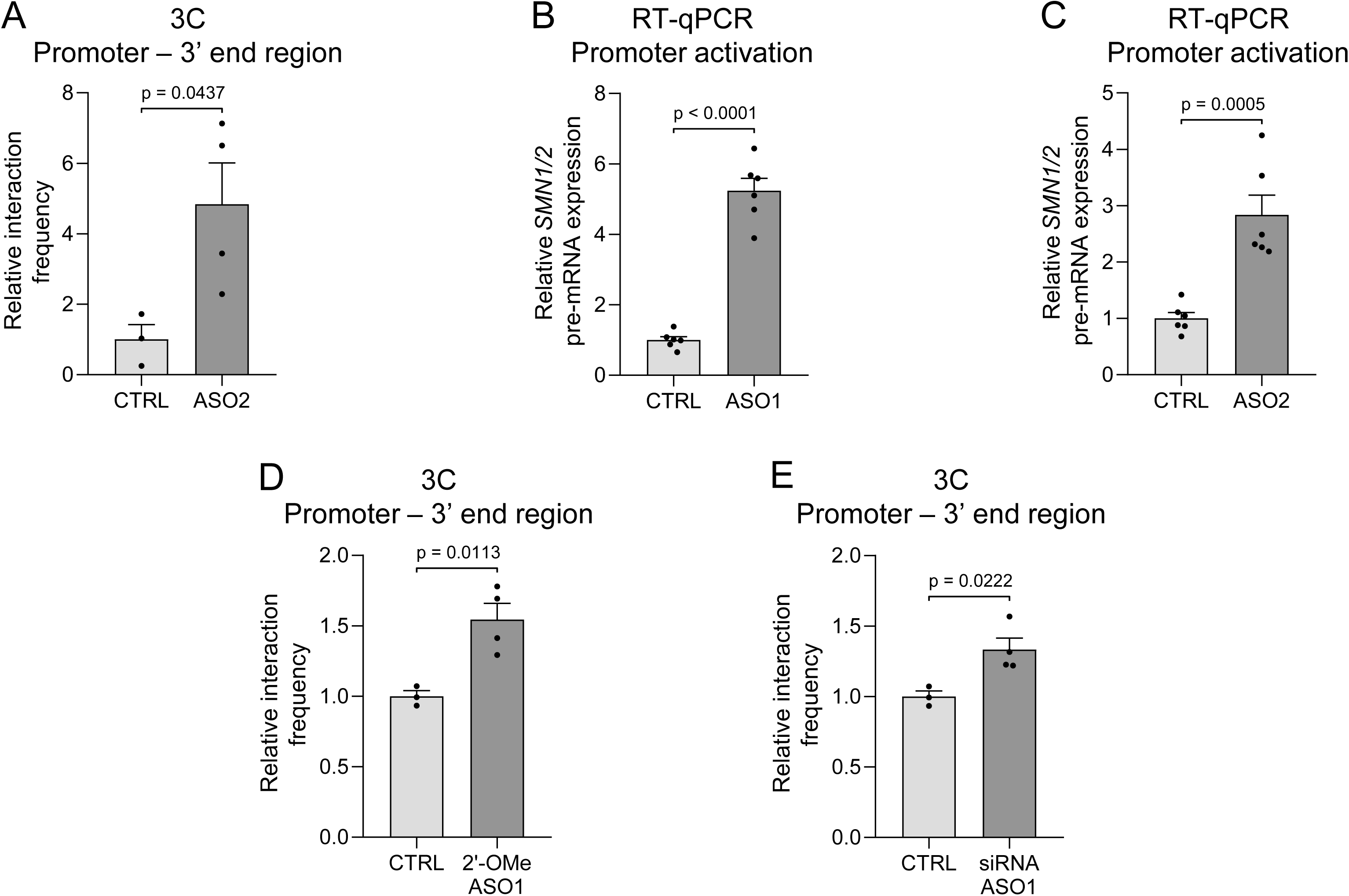
ASO specificity and promoter activation. (A, D–E) 3C-qPCR analysis of *SMN1/2* promoter–3′-end interactions in HEK293T cells transfected for 72 h with or without ASO2 (A), 2′-O-methyl-modified ASO1 (2′-OMe ASO1; D), or a Stealth siRNA (Invitrogen) whose guide strand corresponds to ASO1 (E), all at 100 nM. Interactions were normalized to a U6 gene amplicon and to the corresponding control. n = 3–4 technical replicates; experiments were independently repeated with consistent outcomes. The control in (E) is the same as in (D). (B–C) RT-qPCR of *SMN1/2* promoter activation in cells with or without ASO1 (B) or ASO2 (C) (100 nM, 72 h). E1–I1 pre-mRNA levels were normalized to GAPDH and to the corresponding control. n = 6 biological replicates. Data are mean ± SEM. Two-tailed Student’s *t* test; significant p values are shown; p < 0.05.

One of the possible consequences of the promotion of gene looping by ASOs mapping at the 3’ end of a gene is that they may affect promoter function. Indeed, we found that both ASO1 (Figure 5B) and ASO2 (Figure 5C) upregulate promoter activity.

### Controls on ASO specificity and chemistry

A series of controls were performed to better understand the promotion of looping by ASOs. First, the looping effect does not seem to be related to ASO1’s specific sequence since it is also observed with ASO2 (Figure 5A).

As well as nusinersen, the ASO1 used in this report bears the 2’-O-methoxyethyl (2’-MOE) sugar modification in which the hydroxyl group on the second carbon of every ribose sugar in the oligonucleotide is replaced with a bulky, methoxyethyl group. Figure 5D shows that an ASO with a different chemistry, the 2’-O-methyl (2’-OMe) modification, that upregulates E7 inclusion (Figure S3A), also promotes gene looping.

Finally, since short interfering RNAs (siRNAs) are widely used in therapeutic studies^27^ and were shown to affect chromatin structure^8^, we tested a Stealth (Invitrogen technology) siRNA whose guide strand has the ASO1 sequence and found that it also promotes gene looping (Figure 5E). Interestingly, this siRNA is not as potent as ASO1 to promote E7 inclusion (Figures S3B and C).

### ASO-elicited looping depends on cohesin and CTCF

We next wanted to verify if the ASO-promoted looping was mediated by cohesin. This protein is a key component of long-distance interactions between TAD boundaries and between enhancers and promoters^28^. Typical experiments to assess roles of cohesin involve depletion of its RAD21 subunit. However, Rhodes et al.^29^ and Liu and Dekker^30^ independently showed that depletion of RAD21 in cycling cells caused an accumulation of mitotic cells compared to the control, most probably due to the fact that cohesin also plays an essential role in sister chromatid cohesion and mitotic progression^31^. Since our results were obtained in cycling cells (HEK293T), in order to avoid indirect effects related to the cell cycle, we decided to knock down, via RNAi, the cohesin loader subunit NIPBL1, whose degradation was shown to affect looping but to preserve the proliferative capacity of cycling cells^32^. NIPBL1 knockdown (Figure S4A) abolishes the upregulation of contacts between the promoter and 3’ end regions caused by ASO1 (Figure 6A). Consistently, NIPBL knockdown reduces promoter activation by ASO1 (Figure 6B) which clearly shows that looping is necessary for activation at distance. Most relevant, NIPBL1 knockdown does not inhibit either basal or ASO1-stimulated E7 inclusion (Figure 6C). This clearly indicates that promotion of looping and regulation of alternative splicing are two independent mechanisms caused by ASO1.

**Figure 6.**
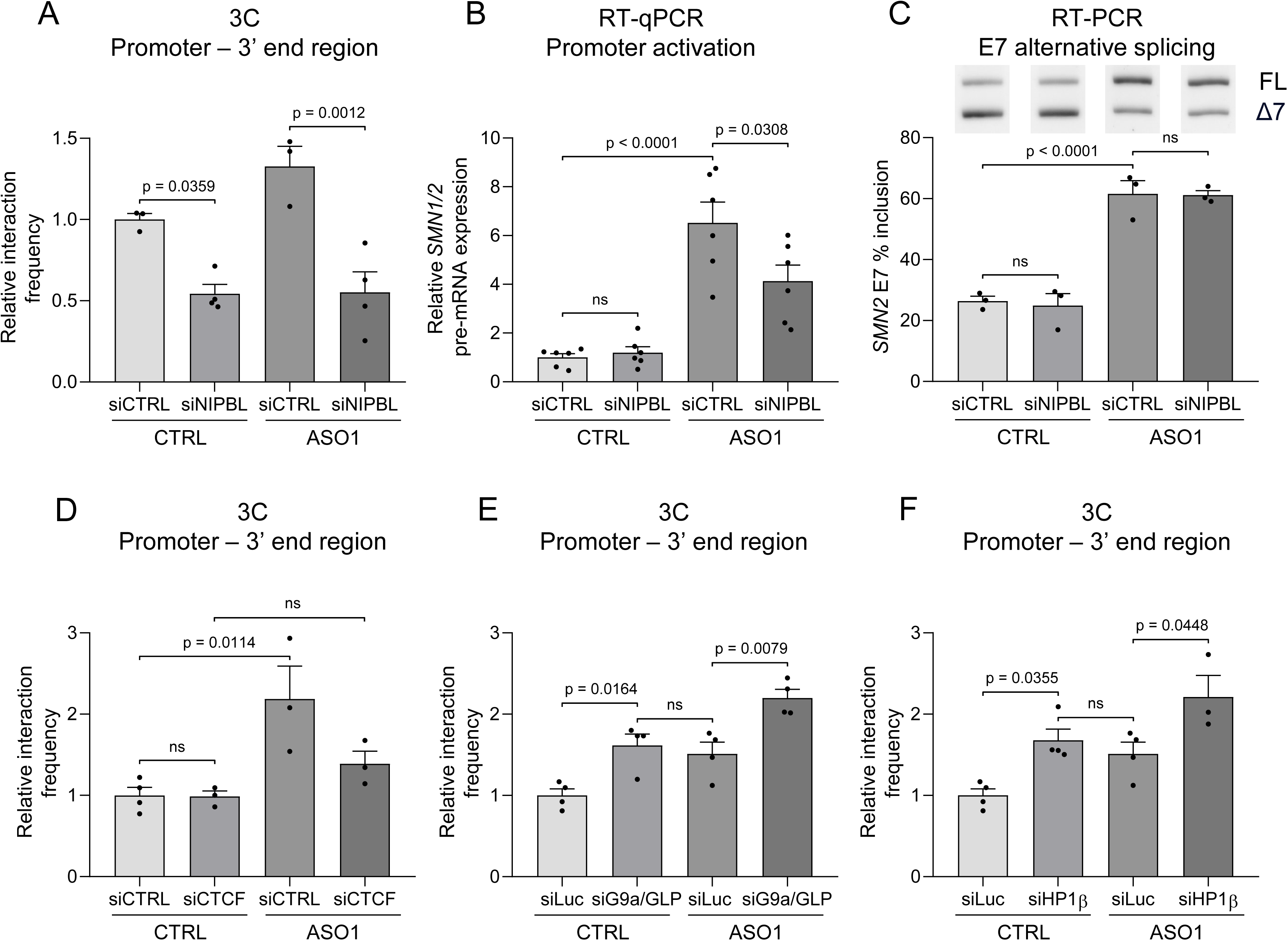
Cohesin and CTCF, but not H3K9 dimethylation, are necessary for the ASO1 promoted gene looping. HEK293T cells were transfected with the indicated siRNAs (50 nM, 72 h), with or without ASO1 (100 nM, 72 h). (A, D–F) 3C-qPCR of *SMN1/2* promoter–3′-end interactions following depletion of NIPBL (A; cohesin-loading factor; siCTRL/siNIPBL), CTCF (D; siCTRL/siCTCF), G9a/GLP (E; siLuc/siG9a/GLP combined knock down), or HP1β (F; siLuc/siHP1β). A and D were normalized to a U6 gene amplicon and siCTRL without ASO1; E and F to a GAPDH gene amplicon and siLuc without ASO1. n = 3–4 technical replicates (A, D, F) and n = 4 (E). A, E, and F were independently repeated with consistent outcomes. The siLuc controls in (F) are the same as in (E). (B–C) siCTRL or siNIPBL cells. (B) RT-qPCR of *SMN1/2* promoter activation; E1–I1 pre-mRNA was normalized to GAPDH and siCTRL without ASO1. n = 6 biological replicates. (C) RT-PCR of *SMN2* E7 splicing; FL, full-length (upper band); Δ7, E7-skipping (lower band); representative gel (top) and E7 inclusion (%) from n = 3 biological replicates (bottom). Data are mean ± SEM. Statistics: two-way ANOVA/Tukey (A, B, D–F) or two-factor beta regression/Tukey (C). Significant p values are shown; ns, not significant; p < 0.05.

The DNA-binding protein CTCF (CCCTC-binding factor) is located at loop boundaries and is required for cohesin accumulation at these sites^33^. The prevailing model is that cohesin packages the chromatin fiber while CTCF defines focal boundaries by constraining packaging^34^. Figure 6D shows that, as well as NIPBL depletion, CTCF knockdown (Figure S4B) reduces looping promoted by ASO1. This is consistent with the finding of CTCF sites at both ends of the SMN genes^35^. It is worth noting that, NIPBL knockdown reduces promoter-3’-end contacts in the absence of ASO1 (Figure 6A) and promotions of looping by ASO1 and targeted acetylation at the E7 region are additive (Figure S5)

### Looping is not caused by H3K9 dimethylation

In view of results in Figure 1E, we speculated that H3K9 dimethylation elicted by ASO1 both at its target site and at the promoter would indirectly promote gene looping. However, experiments in Figures 6E and 6F demonstrated that this is not the case. Depletions of either the methylation enzyme components (Figure 6E) or the H3K9me2 reading factor HP1• (Figure 6F), far from inhibiting promoter-3’ end contacts actually promote them. Controls of RNAi depletion are shown in Figure S1.

### Messenger RNA 3•-end formation contributes to looping

Previous studies in yeast showed that actively transcribed genes can adopt a looped conformation that juxtaposes their promoter and mRNA 3’-end formation regions and that this topology contributes to transcriptional memory and rapid gene reactivation^36,37^. These observations prompted us to determine whether the ASO1-induced gene looping was linked to events occurring at the cleavage and polyadenylation site. We explored two independent experimental approaches. On the one hand, we found that RNAi depletion of three key factors of the cleavage and polyadenylation complex (CPSF73, CstF64 and PCF11, Figure S6) reduces the ASO1-elicited looping (Figure 7A). On the other hand, we replaced in HCT116 cells the endogenous *SMN1/2* exon 8 poly(A) signal and adjacent downstream sequence with an artificial sequence-engineered A-rich (or A75) tract, followed by a self-cleaving ribozyme. This arrangement generates the polyadenylated transcript independently of poly(A)-site recognition and canonical pre-mRNA cleavage. We first confirmed that ASO1 enhanced looping in WT HCT116 cells, recapitulating the effect previously observed in HEK293T cells. This response was attenuated with the polyA (pA) mutant (Figure 7B). Thus, although canonical cleavage and polyadenylation are not strictly required for promoter–3’end communication, they appear to facilitate the full looping response induced by ASO1, consistent with a role for last exon-associated processing events in stabilizing *SMN1/2* gene topology.

**Figure 7.**
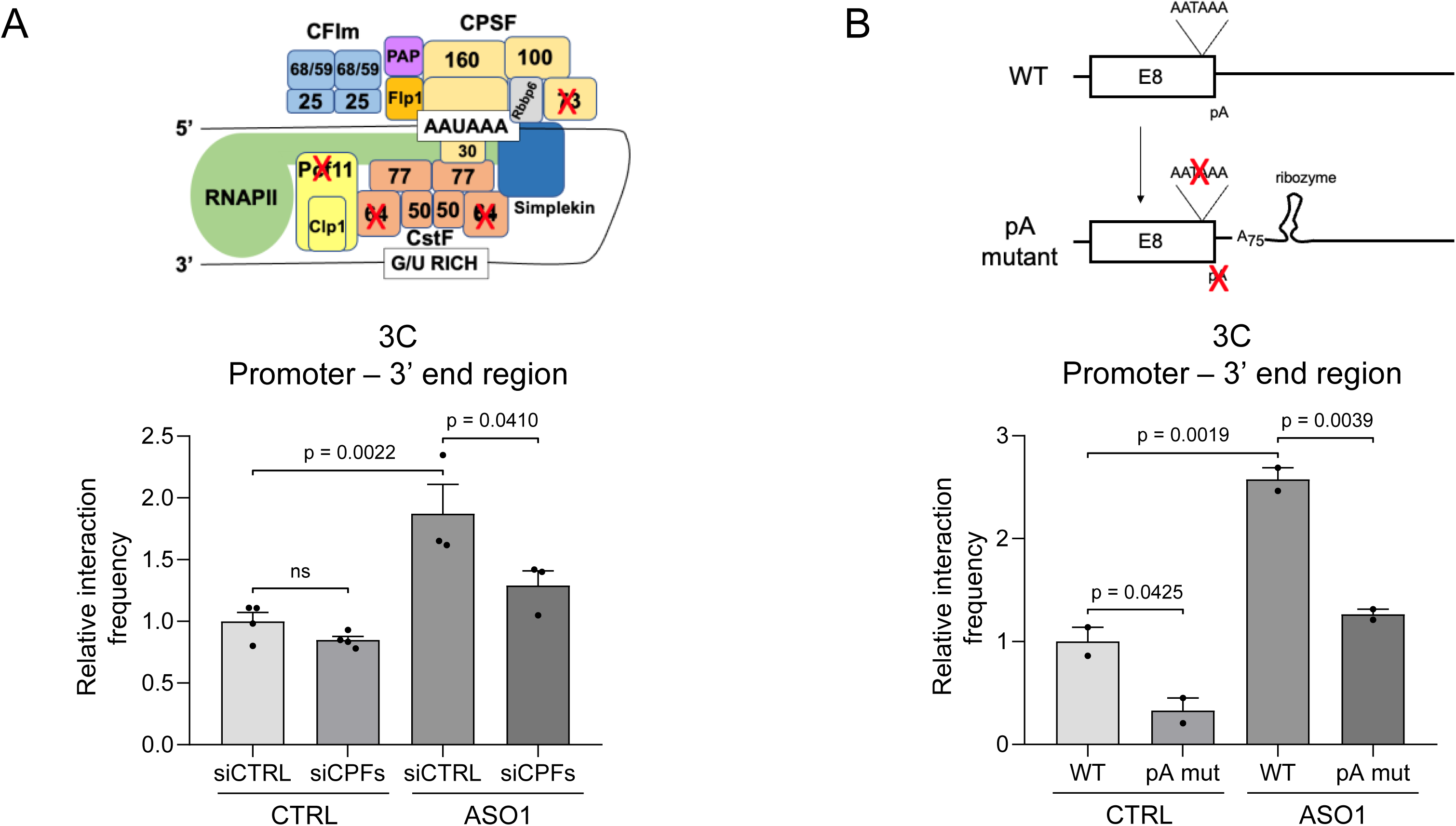
Trans-acting factors and cis-acting sequences of mRNA 3’ cleavage/polyadenylation participate in ASO-elicited gene looping. (A) Top, schematic representation of pre-mRNA 3′-end cleavage/polyadenylation factors, adapted from Davis and Shi^48^. Bottom, 3C-qPCR of *SMN1/2* promoter–3′-end interactions in HEK293T cells transfected with siCTRL or combined siRNAs against CPSF73, CstF64, and PCF11 (siCPFs; 50 nM, 72 h), with or without ASO1 (100 nM, 72 h). Interactions were normalized to a GAPDH gene amplicon and siCTRL without ASO1. n = 3–4 technical replicates. (B) Top, schematic representation of the *SMN1/2* 3′-end region in WT and poly(A)-signal mutant (pA mut) HCT116 cells. In pA mut cells, the endogenous exon 8 poly(A) signal and adjacent downstream sequence were replaced by a synthetic A-rich (A75) tract and self-cleaving ribozyme, generating the transcript 3′ end independently of canonical poly(A)-site recognition and pre-mRNA cleavage. Bottom, 3C-qPCR of promoter–3′-end interactions in WT and pA mut cells, with or without ASO1 (100 nM, 72 h), normalized to a U6 gene amplicon and WT without ASO1. n = 2 technical replicates; independently repeated with consistent outcomes. Data are mean ± SEM. Two-way ANOVA followed by Tukey’s multiple-comparisons test; significant p values are shown; ns, not significant; p < 0.05.

## DISCUSSION

The finding that antisense oligonucleotides used in mRNA splicing correction also cause histone methylation^11,38^ opened a series of questions not only about the underlying mechanisms but also about the gene expression consequences. Accordingly, the antisense oligonucleotide nusinersen (ASO1) displaces the negative splicing factors hnRNPA1/A2 from their target site in the *SMN2* pre-mRNA, which promotes inclusion of the critical exon 7 (E7) into the mature mRNA with clear therapeutic benefit. Also, by promoting local H3K9 dimethylation it has an opposite effect on E7 inclusion. These opposing effects are due to the fact that the E7 has been characterized as a Type II exon, whose inclusion is inhibited by intragenic histone methylation. In a deeper analysis, we show here that knockdown of the histone methylating enzymes G9a and GLP or the H3K9me2 reader HP1•, as well as knockout of G9a alone, mitigate the negative chromatin effects of ASO1 on E7 inclusion (Figures 1B-D). These effects are paralleled by the distribution patterns of the H3K9me mark along the *SMN1/2* merge gene (Figure 1E). A relevant finding was that ASO1 not only promotes histone methylation at its target region, located at the 3’ end of the gene, but also, and distinctively, at the promoter (Figure 1E). This prompted us to further investigate both this and possible distant effects, such as those of targeted histone acetylation.

Targeted histone modifications of chromatin regions associated to alternative splicing events were reported before but not in combination with ASOs^39^. Our experiments of targeted histone acetylation using the dCas9 strategy had a dual purpose. On the one hand, since promotion of global histone acetylation with histone deacetylase inhibitors such as valproic acid (VPA) or trichostatin A (TSA) counteracted the negative effects of histone methylation on E7 inclusion^11^, it was important to rule out an indirect mechanism. On the other hand, we wondered if targeted histone acetylation at the E7 gene region also had effects at the 30-kbp-distant promoter. For the first purpose, results in Figures 2B-D clearly show that targeting dCas9-VP64 to the E7 gene region enhances the ASO1-mediated promotion of E7 inclusion into its mature mRNA and that this effect is due to the ability of VP64 to induce histone acetylation. Moreover, specific histone acetylation at the E7 gene region also reverts the negative effect of ASO2 on E7 inclusion into the mRNA (Figure 2E), which strongly supports the model in Figure 1A, using now a gene-specific histone acetylating tool. These results may have important implications for future antisense therapies in which both ASOs and targeted chromatin-modifying enzymes could be combined in a gene-specific manner, minimizing potential HDACi pleiotropic effects. As far as the second purpose is concerned, we found that targeting dCas9-VP64 to the E7 gene region not only promotes local histone acetylation (Figure 3A), as expected, but also at the promoter (Figure 3B), resulting in its activation (Figure 3D). As a positive control, targeting dCas9-VP64 at the promoter activates it, as expected for a typical CRISPRa experiment (Figure 3C). The fact that targeting histone acetylation at the promoter also affects E7 alternative splicing *per se* and enhances ASO1 promotion of E7 inclusion (Figure 3E) constitutes a strong support, using an endogenous gene, of early findings that promoters^19,20,22^, enhancers^23^ and transcription factors^21,40^ control alternative splicing at distance.

### A new property of ASOs

We demonstrate that, apart from their expected chromatin effects at their respective target regions, both ASO1 and E7-targeted-dCas9-VP64 have distant effects at the promoter region. We therefore decided to investigate the existence of physical proximity between the promoter and the 3’-end region of the 30-kbp-long *SMN* gene using chromosome conformation capture (3C). We set the anchor at the promoter and interrogated fragments at the 3’ end of the gene, at intron 1 and at an upstream enhancer element. In the absence of ASO1, the 3C signal was similar in all three interrogated regions (Figure 4A, light grey bars). However, upon ASO1 transfection, contacts between the promoter anchor and the 3’end region were significantly increased (dark grey bars). We anticipated detection of a basal level of looping in the absence of ASO1 that could *per se* justify the peak of promoter H3K9 dimethylation observed in Figure 1E. However, the finding that looping was increased by ASO1 revealed an unexpected and completely new property of ASOs.

Consistent with the distant effects of targeted acetylation on the promoter and on alternative splicing (Figure 3), directing dCas9-VP64 either to the E7 region (Figure 4B) or to the promoter (Figure 4C) upregulates looping.

We interpret that the main biological consequence of the ASO1-promoted gene looping is the activation of the *SMN2* gene promoter, that occurs both with ASO1 and ASO2 (Figure 5B and C). As controls, an siRNA with the ASO1 sequence and an ASO1 with a different chemistry promote gene looping as well (Figure 5E and D). The fact that the siRNA with the ASO1 sequence mimics ASO1 in looping enhancement but not in alternative splicing reinforces the idea that looping promotion and E7 splicing regulation are independent activities of ASO1.

To dig deeper into the underlying mechanism, the *bona fide* nature of the observed enhancement of looping by ASO1 is corroborated by its abolition upon RNAi depletion of the cohesin loader NIPBL1 and of CTCF (Figure 6A and D). The NIPBL depletion experiments reveal that promoter activation (Figure 6B), but not regulation of E7 splicing (Figure 6C), depends on looping. Experiments in Figures 6E and F complete a complex panorama in which ASO1 hast three apparently independent activities: Firstly, promotion of E7 inclusion by displacement of hnRNPA1/A2 from the pre-mRNA; second, promotion of H3K9 dimethylation and third, promotion of looping and subsequent promoter activation. It should be noted that a database search for transcriptional enhancer sequences and characteristic histone marks around the 3’-end the SMN1/2 gene that may explain promoter activation proved to be negative. With respect to looping, evidence in Figures 6A and 7B indicate that it exists in basal conditions but is enhanced by the ASO. Also Figures 6E, 6F and S7 show that looping is stimulated by factors other than the ASO. Additionally, the demonstration that H3K9 dimethylation is not unnecessary for looping (Figures 6E and F) allows speculation that the methylation peak observed at the promoter (Figure 1E) may be the consequence rather than the cause of looping. Its enhancement by depletion of the writers and reader of the H3K9me2 mark (Figures 6E and F) are the consequence of further chromatin opening. Indeed, chromatin opening by the HDACi VPA (Figure S7) and targeted acetylation (Figure 4B and C) increase 5’-3’ looping *per se*.

Over twenty years ago, the Proudfoot and Hampsey labs^41,42^ showed the existence of transcription-dependent gene loops between promoter and polyadenylation sites (PAS) of protein-coding genes in budding yeast. Since then, genome-wide studies have focused on long-distance interacions that defined TADs and overlooked the importance of closer-range interactions within genes, produced by gene looping. However, two recent studies reveal that genome-wide promoter selection defines PAS selection^43,44^, which highlights the importance of gene looping. Totally consistent with these studies, using two independent perturbation approaches, one *in trans* and the other *in cis*, we now show that canonical mRNA 3•-end formation contributes to the ASO-elicited gene looping in the *SMN1/2* gene (Figure 7).

Apart from obvious therapeutic benefit that a splicing correcting ASO at the same time activates transcription of the regulated gene, our results now provide a rational for the observation that Spinraza activates transcription of the *SMN2* gene both in the endogenous gene and in a minigene reporter^45^.

Promoter activation by ASOs may involve RNAPII itself. Gene loops were shown to be formed by physical interaction of transcriptional initiation and cleavage/polyadenylation factors occupying the opposite ends of a gene^46^. Allepuz-Fuster et al.^47^ demonstrated a direct role for RNAPII in gene looping through its Rpb4 subunit and proposed a mechanism of transfer of RNAPII from the end of a gene to its promoter, which enhances transcriptional reinitiation by the general transcription factor TFIIB.

We foresee that the results in this report will open new avenues for the world of ASO therapies. Splicing-correcting ASO design and testing will not only need assessment of effects at the pre-mRNA level, but also parallel assessment at the chromatin level. This will involve not only deployment of histone marks but also changes in chromatin 3D conformation and transcription. In any case, further investigation will be necessary to decipher the detailed mechanisms of the chromatin effects of ASOs.

## LIMITATIONS OF THIS STUDY

- ASO-mediated gene looping was found and characterized in cultured cells. Its existence in a whole organism exceeds the scope of the present report.
- Whether other ASOs whose target sites map at 3’-end regions of other genes also promote looping and promoter activation remains to be investigated. We focused here on the conditions and participants of an unforeseen property of ASOs.

## AUTHOR CONTRIBUTION

J.N.S. and E.H. performed most of the experiments. L.E.M., A.F. and N.J.P. helped with experiments and ideas. A.R.K. supervised the work and wrote the manuscript.

## ACKNOWLEDGMENTS

We thank Reini Luco for the CRISPR-Cas9 expression vectors, Valeria Buggiano and Mariano López Gringauz for technical assistance, and Nicolás Barberón, Juan Cristobal Muñoz, Luciana Gómez Acuña, Mario Rossi and Elias Friman for helpful discussions. This work was supported by joint grants from Familias Atrofia Muscular Espinal (FAME, Argentina) and CureSMA (USA) and grants from the Lounsbery Foundation (USA). A.R.K. acknowledges support from the Universidad de Buenos Aires (UBACYT 20020170100046BA), the Agencia Nacional de Promoción Científica y Tecnológica of Argentina (PICT-2019 862), the former Ministry of Science, Technology and Innovation of Argentina, closed by the Milei administration (CONVE-2023-100766162-APN-MCT), and Laboratorios Gador. A.R.K. is a career investigator, and J.N.S. and E.H. received Ph.D. fellowships from the Consejo Nacional de Investigaciones Científicas y Técnicas of Argentina (CONICET). L.E.M has received ongoing support from Linacre College, Oxford, though a Paul Nurse junior fellowship.

## STAR METHODS

### Resource availability

#### Lead contact

Further information and requests for resources and reagents should be directed to and will be fulfilled by the lead contact, Dr. Alberto R. Kornblihtt.

#### Cell lines

HEK293T (WT and G9a KO) cell lines were maintained at 37 °C in a humidified incubator with 5 % CO• in DMEM (4.5 g/L glucose; Gibco) supplemented with 10 % fetal bovine serum (FBS) and penicillin/streptomycin (Pen/Strep). The HEK293T CRISPR G9a knockout cell line was previously described^13^ and loss of G9a expression was confirmed in this study by Western blot. Wild-type HEK293T and G9a knockout cells were seeded at equivalent densities and matched for passage number to ensure comparability across experiments.

#### Materials availability

All unique/stable reagents generated in this study are available from the lead contact upon request.

### Method details

#### Cell culture and treatments

HEK293T cells (wild-type and G9a knockout) and HCT116 cells were maintained in DMEM (Gibco) supplemented with 10% FBS and 1% penicillin/streptomycin at 37 °C and 5% CO•. Transfections with plasmids, siRNAs, or ASOs were performed using Lipofectamine 2000 (Thermo Fisher Scientific) in Opti-MEM (Gibco) following the manufacturer’s recommendations, and medium was replaced after 24 h.

For dCas9-VP64 experiments, 1.5 × 10• cells were plated per well in 12-well plates and transfected the next day with 2 µg DNA at a 6:1 molar ratio of LentiGuide-Puro vectors (Addgene #52963), previously cloned to contain the gRNA of interest, to dCas9-VP64-GFP (Addgene #61422). ASOs were typically introduced 48 h after seeding at the indicated concentrations. CBP30 (MedChemExpress, Cat# HY-15826) was tested in parallel at 40 µM as a novel treatment. Cells were collected 72 h after ASO or 96 h after plasmid transfection for RNA, protein, or chromatin extraction. siRNA-mediated knockdown experiments are described separately (see RNAi knockdown section). In experiments without plasmid transfection, ASOs (ASO1, ASO2, 2’-OMe ASO1 or siRNA ASO1) were transfected 24 h post-seeding, and cells were collected 72 h later under the same schedule.

For each figure, one representative experiment with biological replicates is shown, and all experiments were independently repeated more than once to ensure reproducibility.

#### Antisense oligonucleotide synthesis

ASO1 (5•-ATTCACTTTCATAATGCTGG-3•), a seven-mismatch control (5•-AATCATTTGCTTCATACAGG-3•), and ASO2 (5•-AAAGTATGTTTCTTCCACAC-3•) are 2•-O-methoxyethyl-modified oligonucleotides with phosphorothioate backbone and all 5-methyl cytosines. 2’-OMe ASO1 has the same sequence as ASO1 but is 2’-O-methyl-modified with phosphorothioate backbone. All these antisense oligonucleotides were purchased from IDT. They were dissolved in H2O_mq_ and synthesized following the same chemical design strategy previously described^11^.

#### RNA extraction and RT-PCR

Total RNA was isolated using TRIzol reagent (Invitrogen) following the manufacturer’s instructions. One microgram of RNA was reverse-transcribed with M-MLV reverse transcriptase (Thermo Fisher Scientific) and oligo-dT primers, and the resulting cDNA was amplified. Amplification and analysis of *SMN2* transcripts were performed as described^12^.

For PCR amplification, T-Plus Free DNA Polymerase (Inbio Highway, Cat# K1009) was used with *SMN2* primers surrounding exon 7 (listed in Table S1). PCR products were resolved on 6% acrylamide (29:1) TBE gels and stained with ethidium bromide. Bands were visualized using a UVP Gel Imaging and Documentation System, quantified with ImageJ.

#### Promoter activation assays (RT–qPCR)

Total RNA was extracted as described above and reverse-transcribed with random primers. For each sample, parallel +RT and −RT reactions were prepared to monitor genomic DNA carryover; samples with high −RT signal at the target amplicon were excluded. Pre-mRNA levels were quantified by SYBR Green qPCR using primers spanning the *SMN2* exon 1–intron 1 junction (*SMN2* E1–I1; sequences in Table S1). Absolute/relative quantities were obtained from a standard curve generated from a serial dilution (pooled cDNA from experimental samples) run on every plate. Target quantities were normalized to the reference gene indicated in the figure legends (primers in Table S1) and compared relative to the control condition unless otherwise noted.

#### Western blot

Cells were lysed in 2× Laemmli buffer (4% SDS, 10% 2-mercaptoethanol, 20% glycerol, 0.004% bromophenol blue, 0.125 M Tris-HCl pH 6.8) and incubated at 95 °C for 5–10 min. Protein samples were separated on 8–15% SDS-PAGE gels (Bio-Rad) and electroblotted onto nitrocellulose membranes. Membranes were blocked in 5% milk in TBS-T, probed with primary antibodies, and incubated with secondary antibodies following the manufacturers’ recommendations. Fluorescent signals were acquired using the LI-COR Odyssey 9120 imaging system, and band intensities were quantified with Image Studio Lite software (LI-COR).

The following antibodies were used:

- Mouse monoclonal anti-H3K9me2 (Abcam, ab1220)
- Rabbit polyclonal anti-H3K9ac (Abcam, ab4441)
- Rabbit polyclonal anti-Histone H3 (acetyl K9 + K14 + K18 + K23 + K27) (Abcam, ab47915)
- Rabbit polyclonal anti-H3K27ac (Abcam, ab4729)
- Rabbit polyclonal anti-H3 (Millipore, 07-690)
- IRDye 800CW Goat anti-Mouse IgG (LI-COR, 926-32210)
- IRDye 800CW Goat anti-Rabbit IgG (LI-COR, 926-32211)

#### RNAi knockdown

Downregulation of G9a (EHMT2), GLP (EHMT1), HP1• (CBX1), NIPBL, CTCF, CPSF73, CstF64, and PCF11 was performed using ON-TARGETplus SMARTpool siRNA oligonucleotides (Dharmacon) at a final concentration of 50 nM. siRNA oligos were transfected into cells following the manufacturer’s instructions and incubated for 72 h to allow efficient knockdown. As a negative control, a non-targeting siRNA control pool (Dharmacon, D-001810-10-20) or siLuc (Dharmacon, P-002099-01-20) was used. Knockdown efficiency was validated by qPCR using the corresponding primers listed in Table S1.

#### Chromatin immunoprecipitation (ChIP) followed by qPCR

Chromatin immunoprecipitation was performed using HEK293T cells, approximately 1 × 10• cells corresponding to a confluent 10 cm culture dish per sample. Cells were crosslinked with 0.5% (v/v) formaldehyde for 10 min at room temperature, and the reaction was quenched with 125 mM glycine. Cells were washed twice with cold PBS and subjected to a two-step lysis procedure: first with a cytoplasmic lysis buffer to isolate nuclei, followed by nuclear lysis buffer to release chromatin.

Chromatin was sheared to an average size of 200–500 bp by sonication (Bioruptor, Diagenode) and diluted in immunoprecipitation (IP) buffer (15 mM Tris-HCl pH 8, 150 mM NaCl, 1 mM EDTA, 1% Triton X-100, 0.01% SDS, and protease inhibitor cocktail; Sigma-Aldrich, Cat# 05056489001). For each IP, 8 •g of chromatin were incubated with 1 •g of antibody or IgG control, together with pre-blocked Dynabeads Protein G (Invitrogen), overnight at 4 °C. The following antibodies were used: rabbit polyclonal anti-H3K9me2 (Abcam, ab1220), goat polyclonal anti-H3 (Abcam, ab12079), rabbit polyclonal anti-H3K9ac (Abcam, ab4441), and rabbit polyclonal anti-Histone H3 (acetyl K9 + K14 + K18 + K23 + K27) (Abcam, ab47915). Beads were washed sequentially with low-salt buffer (20 mM Tris-HCl pH 8, 150 mM NaCl, 2 mM EDTA, 1% Triton X-100, 0.1% SDS), high-salt buffer (20 mM Tris-HCl pH 8, 500 mM NaCl, 2 mM EDTA, 1% Triton X-100, 0.1% SDS), LiCl buffer (10 mM Tris-HCl pH 8.0, 1 mM EDTA, 250 mM LiCl, 1% NP-40, 1% sodium deoxycholate), and once with TE buffer (10 mM Tris-HCl pH 8.0, 1 mM EDTA). Chromatin was eluted in 1% SDS and 100 mM NaHCO• buffer, and crosslinks were reversed overnight at 65 °C.

Recovered DNA was purified using the QIAquick PCR Purification Kit (Qiagen) according to the manufacturer’s instructions. DNA was quantified with a Qubit fluorometer, and enrichment was analyzed by SYBR Green real-time qPCR. A standard curve was included for normalization by linear regression. Primer sequences are listed in Table S1.

#### Guide RNA design

Guide RNAs (gRNAs) were designed for use with the dCas9 system employing the LentiGuide-Puro vector (Addgene plasmid #52963) following the Zhang laboratory two-vector framework^49,50^. Candidate target sequences were identified using the UCSC Genome Browser and evaluated with the CRISPR-Cas9 gRNA Design Checker (Integrated DNA Technologies, IDT) to maximize on-target efficiency and minimize potential off-target effects.

In addition, a non-targeting control gRNA (5•-GCACTCACATCGCTACATCA-3•), designed with no predicted binding sites in the human genome, was generated. Its sequence was validated using NCBI BLAST to ensure absence of off-target alignments. This control was included in all experiments to account for non-specific dCas9 effects. All gRNA sequences used in this study are provided in Table S1.

#### Guide RNA cloning and vector construction

gRNAs were cloned into LentiGuide-Puro following the protocol of Sanjana et al.^49^ and Shalem et al.^50^. The vector was digested overnight with BsmBI and dephosphorylated with FastAP (Thermo Fisher Scientific). The ∼8.3 kb backbone was gel-purified using the QIAquick Gel Extraction Kit (Qiagen).

Annealed oligonucleotides (IDT) were ligated into the digested vector using T4 DNA Ligase (Promega) and transformed into NEB Stable chemically competent E. coli (New England Biolabs). Colonies were selected and verified by colony PCR and Sanger sequencing (Macrogen Inc.). This approach yielded both the target-specific gRNA plasmids and the validated non-targeting control construct used throughout this work.

#### Chromosome Conformation Capture (3C)

3C assays were performed in HEK293T cells following established protocols^24,51^ . Approximately 1 × 10• cells per condition (10 cm confluent dish, ∼1.8 × 10• cells seeded) were used. Depending on the experimental design, cells were transfected with combinations of plasmids, ASOs, or siRNAs. For dCas9-mediated looping experiments, cells were transfected 24 h post-seeding with 15 µg LentiGuide-Puro (Addgene #52963) containing the gRNA of interest and 5 µg dCas9-VP64-GFP (Addgene #61422). ASO1 (100 nM final concentration) was introduced 24 h later, either alone or in combination with SMARTpool siRNAs (Dharmacon; 50 nM) for knockdown experiments. Cells were harvested 72 h after ASO transfection (96 h post-plasmid transfection).

Cells were trypsinized (0.25% trypsin, 37 °C, 5 min), resuspended in DMEM with 10% FBS, pelleted (400 × g, 3 min), and washed in PBS + 10% FBS (500 µl per 1 × 10• cells). Crosslinking was performed with 0.5% formaldehyde (10 min, room temperature), quenched with 0.125 M glycine (5 min, on ice), and centrifuged (225 × g, 8 min, 4 °C). Pellets were lysed in cold buffer (10 mM Tris-HCl pH 7.5, 10 mM NaCl, 5 mM MgCl•, 0.1 mM EGTA, protease inhibitors), incubated on ice (10 min), and nuclei were pelleted (400 × g, 5 min, 4 °C). Nuclei were incubated in 1.2× CutSmart buffer with 0.3% SDS (37 °C, 1 h, 900 rpm), neutralized with 2% Triton X-100 (37 °C, 1 h, 900 rpm), and digested overnight with 500 U of Hinf1 (New England Biolabs).

Following digestion, SDS was inactivated (0.3%, 65 °C, 25 min, 900 rpm), and ligation was carried out in 1.15× ligation buffer containing 2% Triton X-100 and 100 U T4 DNA ligase (New England Biolabs) (16 °C, 4 h, followed by 30 min at room temperature). Crosslinks were reversed with proteinase K (10 mg/ml, 65 °C, overnight), samples were treated with RNase A (10 mg/ml, 37 °C, 30–45 min), and DNA was purified by phenol-chloroform extraction and ethanol precipitation (−20 °C, overnight). DNA pellets were centrifuged (2200 × g, 45 min, 4 °C), washed in 70% ethanol, and resuspended in 500 µl of water.

Digestion efficiency was assessed by qPCR comparing undigested and digested samples with primers flanking restriction sites, ensuring that only samples with efficient digestion were carried forward for interaction analysis. Chromatin interaction frequencies were quantified by qPCR using primers specific for targeted genomic regions and normalized to control fragments to allow comparison between conditions.

#### Generation of the *SMN1/2* poly(A) mutant cell line

An *SMN1/2* 3•-end replacement line was generated in HCT116 cells by CRISPR-Cas9-mediated knock-in. Two sgRNAs were used to excise the endogenous exon 8 poly(A) signal and adjacent downstream sequence. This region was replaced with a donor construct containing lacking a poly(A) signal, a sequence-engineered synthetic 75-nt A-rich tract incorporating dispersed nucleotide substitutions to minimize polymerase slippage or abortive transcription, followed by a self-cleaving ribozyme. This configuration generated a genomically encoded poly(A)-like tail while uncoupling transcript 3•-end formation from canonical cleavage and polyadenylation. A loxP-flanked puromycin-resistance cassette located downstream of the engineered sequence was used for clonal selection. Following isolation of puromycin-resistant clones, the selection cassette was removed by transient expression of Cre recombinase. WT and pA-mutant cells were subsequently treated with control ASO or ASO1 as described above. *SMN1/2* 5•–3• proximity was determined by 3C-qPCR using the same primer pairs and normalization procedure employed for the parental cell line.

#### Quantification and statistical analysis

Band intensities from splicing RT-PCR assays were quantified using FIJI (ImageJ). Raw band intensity (Io) was corrected for local background (Bg) to obtain corrected intensity values (Ic = Io − Bg). Exon 7 inclusion was calculated as:

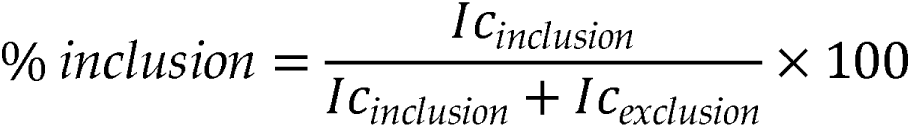

Because of the high sequence similarity between *SMN1* and *SMN2*, PCR amplification with exon 7 flanking primers does not distinguish transcripts derived from each gene. To specifically quantify *SMN2*-derived transcripts, PCR products were digested with DdeI, which recognizes a restriction site unique to exon 8 of *SMN2* (CTNAG). This digestion generated two *SMN2*-specific fragments of 382 and 328 bp, corresponding to exon 7 inclusion and exclusion events, respectively. Band intensities from these fragments were measured in ImageJ to determine the percentage of exon 7 inclusion under each experimental condition, as originally described^12^.

Statistical analyses were performed in R (R Core Team, *R: A Language and Environment for Statistical Computing*). Because exon 7 inclusion is a continuous proportion bounded between 0 and 1, splicing data were analyzed using beta regression models. For factorial experiments, models included the two experimental factors and their interaction, and pairwise comparisons were performed using Tukey-adjusted multiple comparisons. Model fit and residual diagnostics for beta regression models were evaluated using the DHARMa package.

For continuous response variables analyzed under a Gaussian framework, comparisons between two groups were performed using two-tailed Student’s *t* tests, whereas experiments involving two experimental factors were analyzed by two-way ANOVA followed by Tukey’s multiple-comparisons test. Normality was evaluated using the Shapiro–Wilk test and inspection of model residuals, and homogeneity of variances was assessed using Levene’s test. Statistical significance was defined as *p* < 0.05.

Graphical representation of the data was generated using GraphPad Prism 8. Unless otherwise indicated, data are presented as mean ± SEM with individual biological replicates shown as points.

## Supplemental items titles

**Table S1. Oligonucleotide primers used in this paper** (related to Methods).

## Legends to supplemental figures

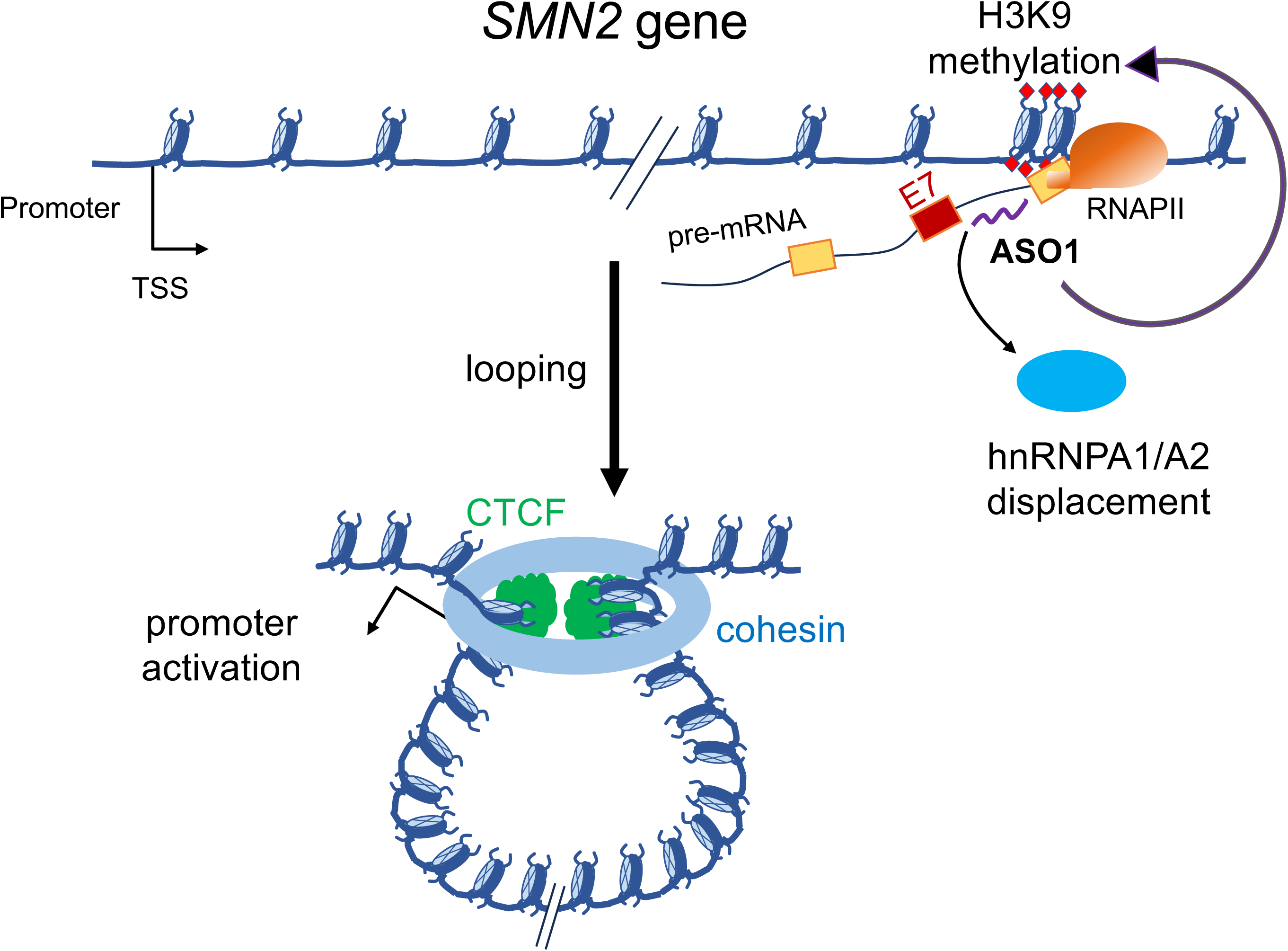
In brief. Three different promoting activities of the splicing-correcting antisense oligonucleotide nusinersen (ASO1 in this report) used in the thearpy of spinal muscular atrophy: Exon 7 inclusion into the SMN2 gene mRNA through base-pairing to sequences in the pre-mRNA and displacement of the negative splicing factors hnRNPA1/A2, deployment H3K9me2 marks (red diamonds), and looping between the promoter and the 3’end region of the gene, with subsequent promoter activation.

**Figure S1.**
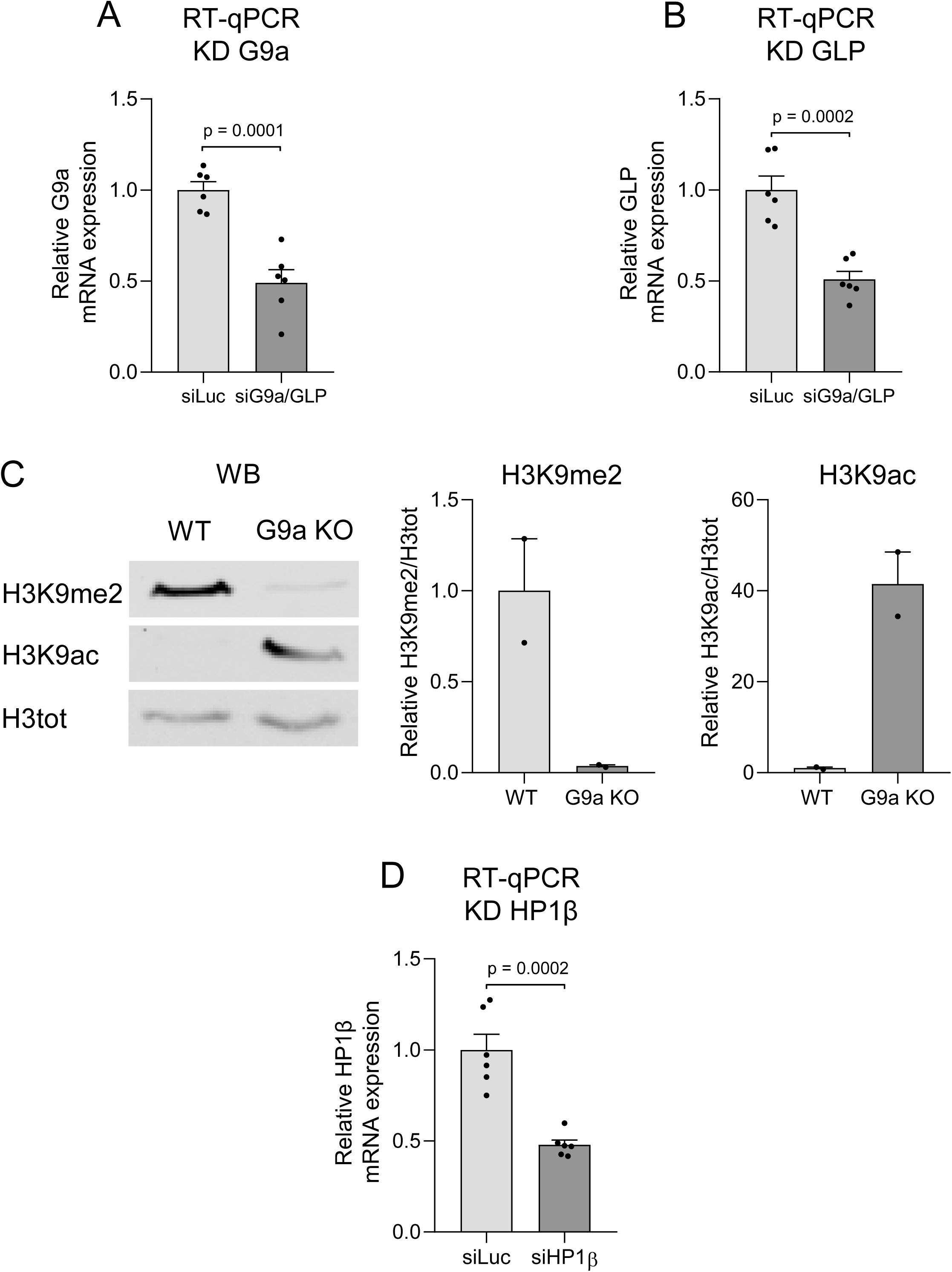
Additional validation of chromatin and knockdown effects (related to. Figure 1) (A–B, D) RT-qPCR validation of G9a/GLP (A–B) and HP1β (D) knockdown in HEK293T cells transfected with siLuc or combined siG9a/siGLP (A–B) or siHP1β (D) (50 nM, 72 h). G9a (A), GLP (B), and HP1β (D) mRNA was normalized to GAPDH and siLuc. n = 6 biological replicates. (C) Western blot of WT and G9a KO HEK293T cells under basal conditions: representative H3K9me2, H3K9ac, and total H3 blots (left), and H3K9me2/H3K9ac quantification normalized to total H3 and WT (right). n = 2 biological replicates. Data are mean ± SEM. A, B, and D, two-tailed Student’s *t* test; significant p values are shown; p < 0.05.

**Figure S2.**
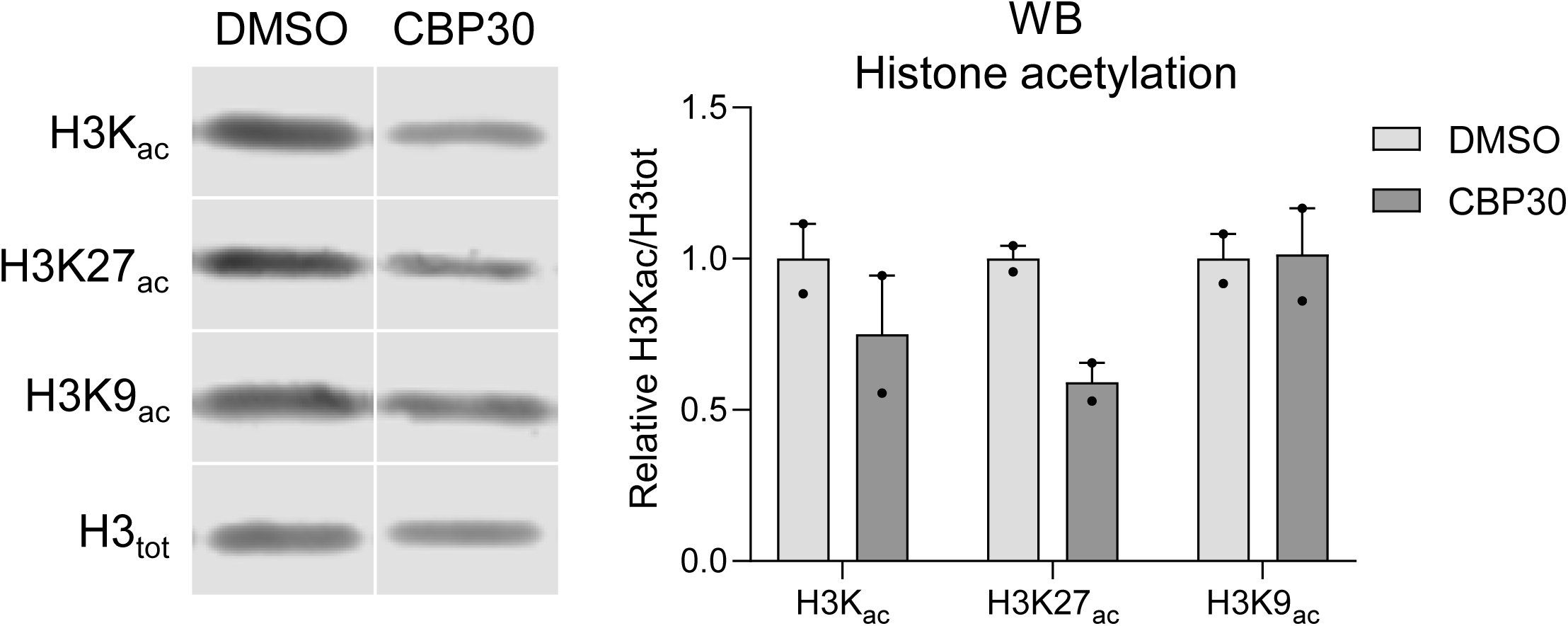
Histone acetylation mediates the synergistic effect between ASO1 and dCas9-VP64 targeting the SMN2 E7 region (related to. Figure 2) Western blot of HEK293T cells treated with DMSO vehicle (0.2%, 48 h) or selective p300/CBP inhibitor CBP30 (40 µM, 48 h). Representative blots (left) and quantification (right) of pan-H3 acetylation (H3Kac; H3K9ac, H3K14ac, H3K18ac, H3K23ac, H3K27ac), H3K27ac, and H3K9ac, normalized to total H3 and DMSO. n = 2 biological replicates; data are mean ± SEM.

**Figure S3.**
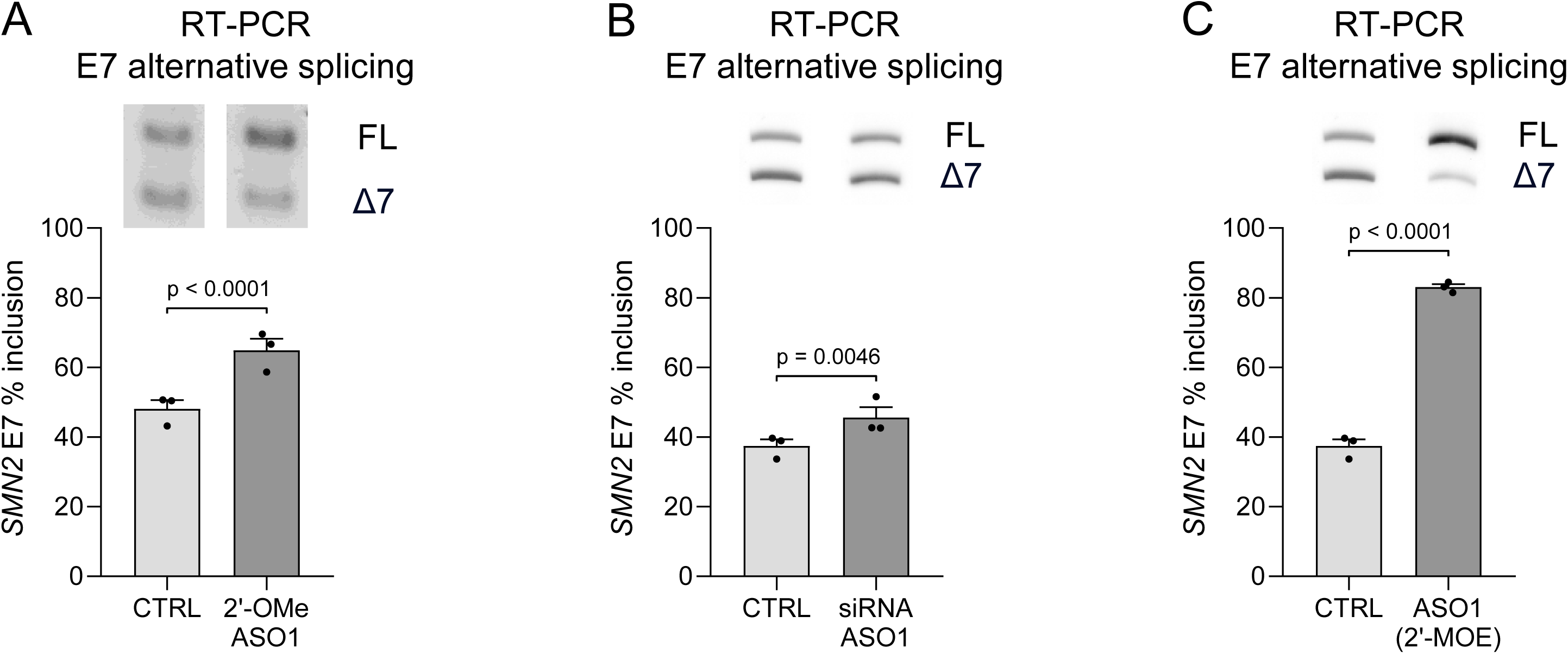
Effects of short nucleic acids other than ASO1 on SMN2 E7 inclusion (related to. Figure 5) (A–C) RT-PCR of *SMN2* E7 splicing in HEK293T cells with or without 2′-OMe ASO1 (A), a Stealth siRNA (Invitrogen) whose guide strand corresponds to ASO1 (B), or standard 2′-MOE ASO1 (C), all at 100 nM for 72 h. FL, full-length (upper band); Δ7, E7-skipping (lower band); representative gels (top) and E7 inclusion (%) from n = 3 biological replicates (bottom). The no-oligonucleotide control in (C) is the same as in (B). Data are mean ± SEM. Beta regression models; significant p values are shown; p < 0.05.

**Figure S4.**
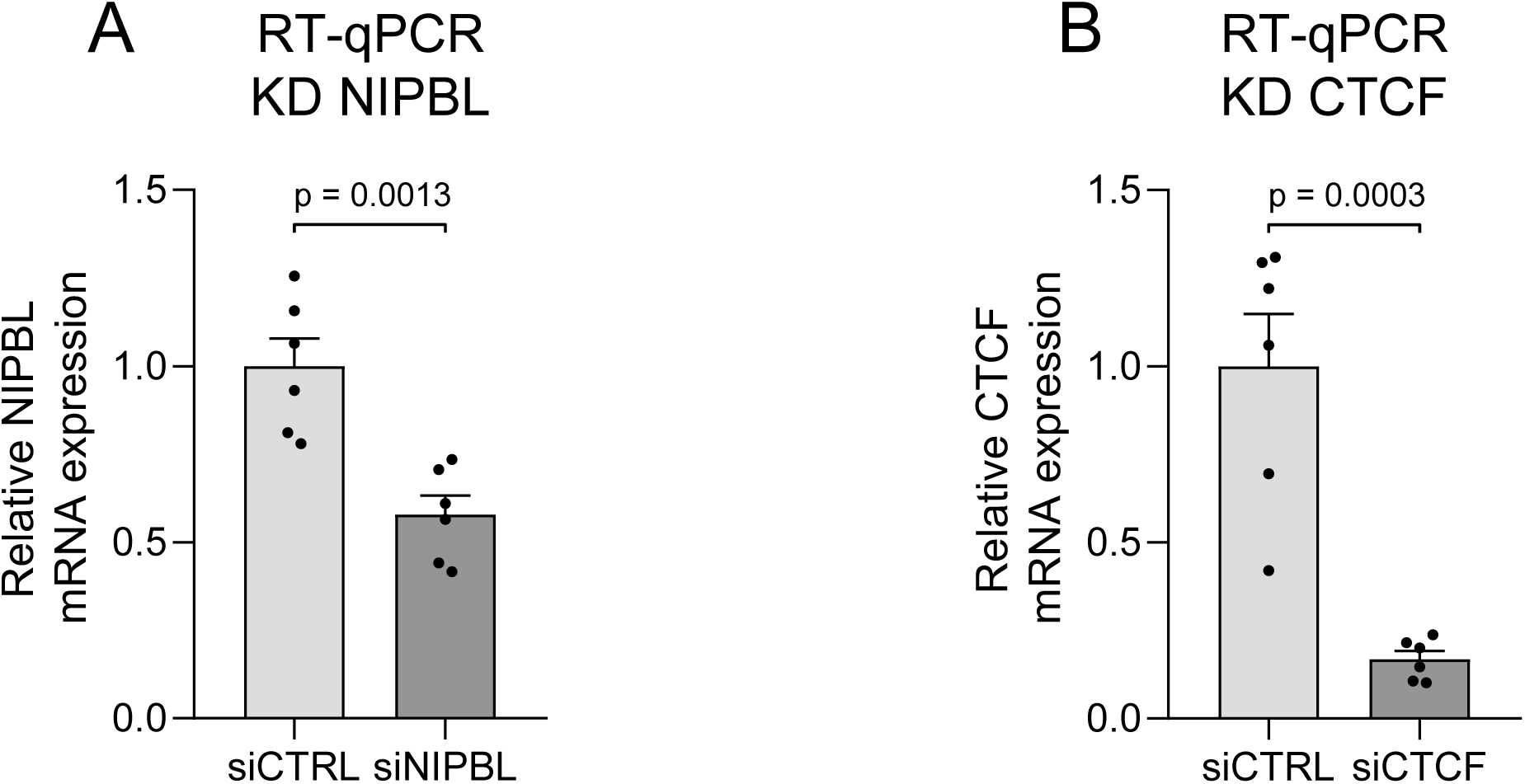
Validation of NIPBL and CTCF knockdowns (related to. Figure 6) (A–B) RT-qPCR validation of NIPBL (A) and CTCF (B) knockdown in HEK293T cells transfected with siCTRL or siNIPBL (A) or siCTCF (B) (50 nM, 72 h). mRNA was normalized to GAPDH and corresponding siCTRL. n = 6 biological replicates; data are mean ± SEM. Two-tailed Student’s *t* test; significant p values are shown; p < 0.05.

**Figure S5.**
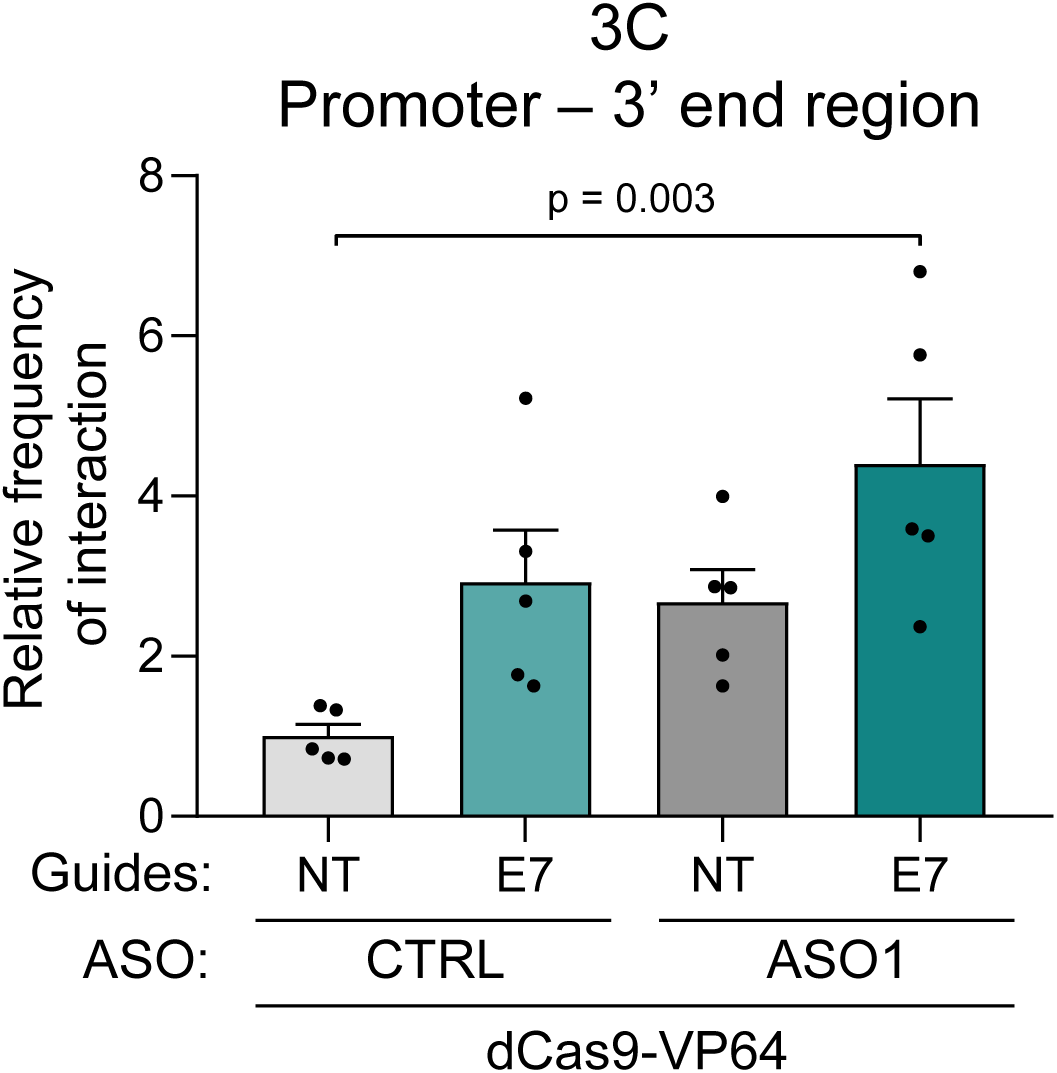
Combined ASO1/dCas9-VP64 targeting enhances promoter–3• interactions (related to. Figure 4) 3C-qPCR of *SMN1/2* promoter–3′-end interactions in HEK293T cells cotransfected with dCas9-VP64 and NT or E7-targeting sgRNAs (96 h), with or without ASO1 (100 nM, 72 h). Interactions were normalized to a U6 gene amplicon and NT without ASO1. n = 5 technical replicates; data are mean ± SEM. Two-way ANOVA/Tukey; significant p values are shown; p < 0.05.

**Figure S6.**
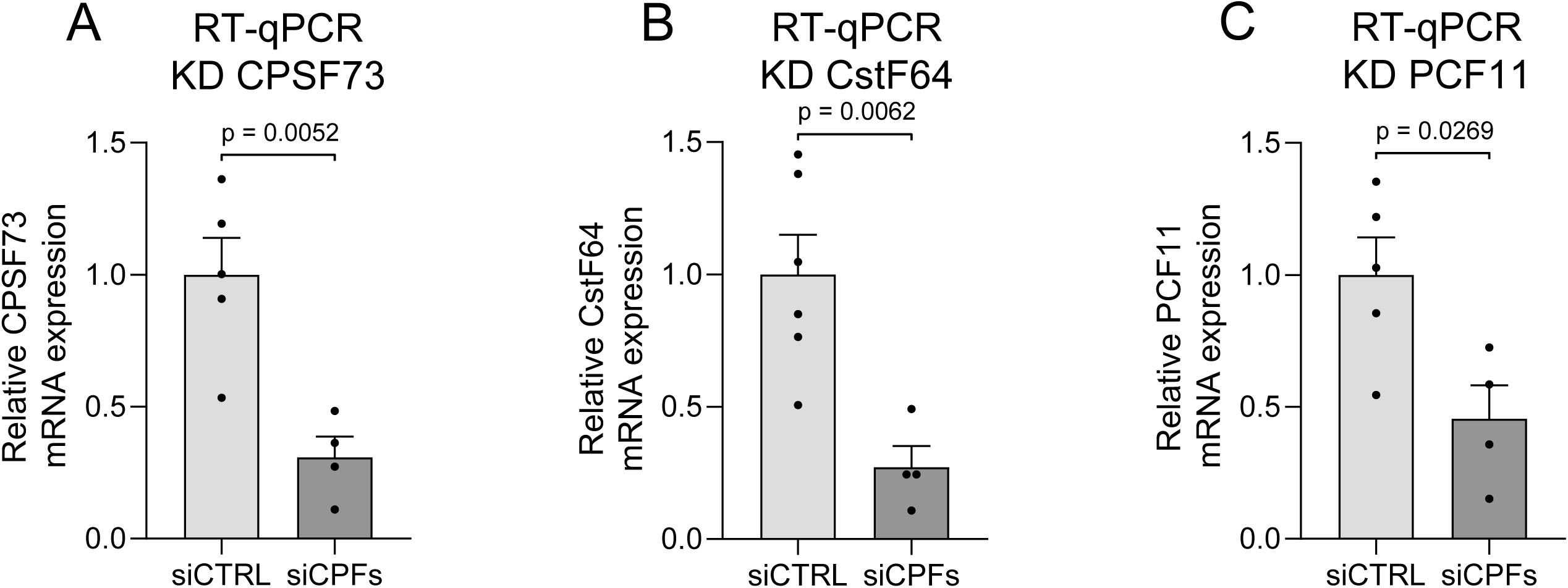
Validation of the knockdown of three different cleavage and polyadenylation factors (related to. Figure 7) (A–C) RT-qPCR validation of CPSF73 (A), CstF64 (B), and PCF11 (C) knockdown in HEK293T cells transfected with siCTRL or combined siRNAs targeting the three cleavage/polyadenylation factors (siCPFs; 50 nM, 72 h). mRNA was normalized to GAPDH and siCTRL. n = 4–6 biological replicates; data are mean ± SEM. Two-tailed Student’s *t* test; significant p values are shown; p < 0.05.

**Figure S7.**
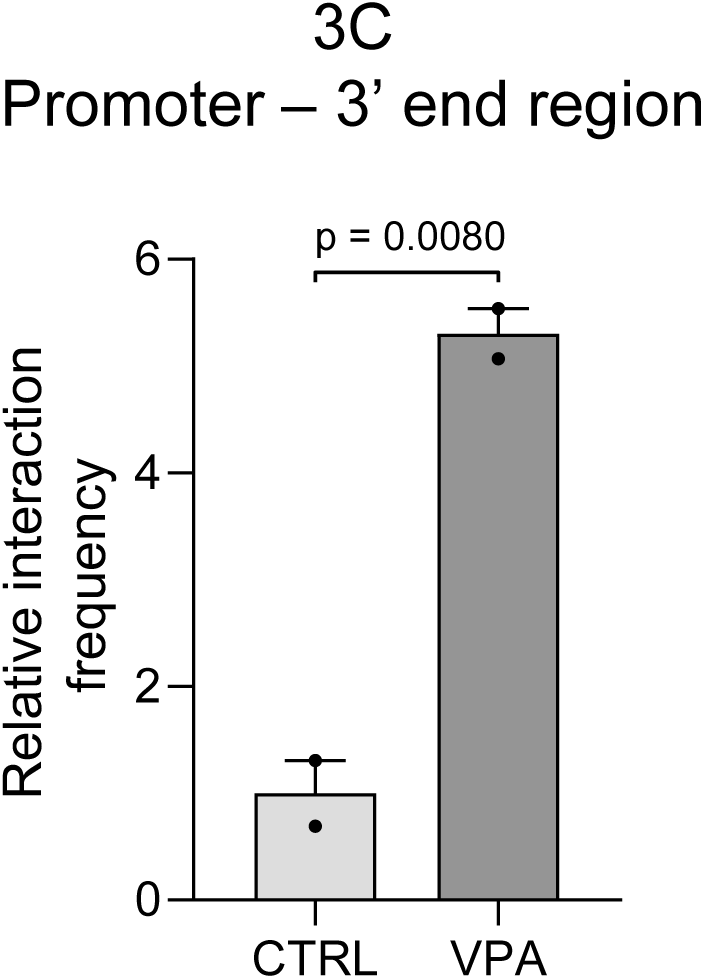
VPA increases looping between the 3’ and 5’ ends of the *SMN1/2* gene (cited in the Discussion) 3C-qPCR of *SMN1/2* promoter–3′-end interactions in HEK293T cells with or without VPA (20 mM, 24 h), normalized to a GAPDH gene amplicon and the control without VPA. n = 2 technical replicates; data are mean ± SEM. Two-tailed Student’s *t* test; significant p values are shown; p < 0.05.

## Notes

### Competing Interest Statement

The authors have declared no competing interest.

### Summary of Updates

The new version contains 11 new experiments shown in New Figures 5C, 5D, 5E, 6B, 6D, 6E, 6F, 7A, 7B and S7, plus new validations and controls shown in New Figures S3A, S3B, S3C, S6A, S6B and S6C, which totals 16 new figures or panels. Summarizing, the revised version adds these new concepts to the ones already shown in the first version: 1. Looping depends not only on cohesin but also on the transcription factor CTCF 2. Looping occurs in basal conditions but is enhanced by ASO1 3. Looping is needed for promoter activation by ASO1 but not for regulation of alternative splicing by ASO1 4. Looping does not seem to depend on the chemistry of the ASO: both 2 prime-MOE and 2 prime-OMe ASO1s cause looping 5. Looping is also stimulated by an siRNA whose guide strand bears the ASO1 sequence 6. Although ASO1 promotes H3K9 dimethylation, this process does not seem to be the cause of looping. On the contrary, the finding of an increase of the H3K9me2 mark at the promoter, 30-kb upstream of the ASO1 target site seems to be the consequence of looping. 7. Perhaps the most significant novelty of the revised version is the finding that ASO-stimulated looping involves the pre-mRNA cleavage and polyadenylation machinery. We show this with two independent experimental approaches: RNAi depletion of key cleavage/polyadenylation factors and CRISPR mutation of the poly(A) signal AATAAA in the endogenous SMN2 gene.

